# A Neural Mass Modelling Framework for Evaluating EEG Source Localisation of Seizure Activity

**DOI:** 10.64898/2026.03.18.712805

**Authors:** Pok Him Siu, Philippa J. Karoly, Sina Mansour L., Artemio Soto-Breceda, Levin Kuhlmann, Mark J. Cook, David B. Grayden

## Abstract

Electroencephalography and magnetoencephalography (EEG/MEG) provide non-invasive measurements of large-scale neural activity but do not directly reveal the underlying cortical sources, motivating the use of source localisation algorithms. However, objective evaluation of these methods remains challenging due to the absence of an experimentally verifiable ground truth. This study presents a simulation framework for generating biologically plausible ictal dynamics and their corresponding EEG signals to enable systematic benchmarking of source imaging approaches. Cortical seizure initiation and propagation were simulated using network-coupled neural mass (Epileptor) models, and combined with realistic forward models of the human head to produce macroscopic, electrophysiological data with known ground truth under varying conditions. Using this dataset, we evaluated established source localisation methods across idealised and realistic scenarios. Existing approaches achieved reasonable spatial accuracy under high-density, noise-free conditions; however, performance degraded substantially with reduced sensor coverage and added noise. This degradation was driven primarily by failures to recover source polarity, even when spatial localisation remained relatively accurate. These results suggest that current methods may be sufficient for identifying epileptogenic regions or tracking regional recruitment, but highlight polarity reconstruction as a key limitation for studies of seizure dynamics and network organisation. The proposed framework provides a reproducible and biologically grounded testbed for the development and evaluation of electrophysiological source localisation techniques.

## 1 Introduction

A pervasive problem with macroscopic, electrophysiological recordings (i.e., EEG/MEG) is that these measurements are a weighted sum of neural activity and, by themselves, do not directly define the underlying sources of neural activity. Implanted stereo-EEG (sEEG) electrodes are able to more directly measure this source activity, but are a highly invasive process and cannot cover the entire brain. Source imaging aims to recreate the neural activity of the underlying sources from the EEG/MEG data using mathematical techniques. However, the underlying problem remains that the sources estimated by these algorithms generally cannot be directly experimentally verified, making it difficult to evaluate and compare the effectiveness of different source imaging algorithms.

By introducing a realistic simulation of some underlying sources of neural activity with their corresponding EEG activity, a test set with a known ground truth can be used to understand the differences between different source localisation methodologies and conditions. Previous studies have typically focused on one to a few sources of various spatial extents (Cioppa et al., 2023; Giri et al., 2022; Hecker et al., 2021; Morik et al., 2024; Petrov, 2012; Rong et al., 2025). However, such simplified source configurations provide limited biological plausibility given that the brain is known to be a highly interconnected and complex hierarchical structure (Burt et al., 2018; Eguíluz et al., 2005; Eickhoff et al., 2018), capable of producing rhythmic and wavelike activity with wavelengths spanning several centimetres (Doelling & Assaneo, 2021; Koller et al., 2024; Pang et al., 2023). Therefore, there remains a need for more adaptable ground-truth simulation frameworks to evaluate source imaging, specific to the application in mind.

Neural mass models are a class of physiologically informed dynamical models, capable of reproducing a diverse range of experimentally observed brain activity (Breakspear, 2017; Siu et al., 2022), such as seizures (Karoly et al., 2018; Ying et al., 2023), sleep (Costa et al., 2016) and brain rhythms (Nunez, 1974; Sotero et al., 2007). Beyond replicating observable features of neural activity, these generative models of brain dynamics provide interpretable parameters that offer insight into the mechanisms that underlie specific brain phenomena Karoly et al., 2018; Maran et al., 2025. Within epilepsy, a widely used example is the Epileptor model, which is a phenomenological dynamical model that has remarkable success in simulating seizure-like activity (V. K. Jirsa et al., 2014). The key distinction (and hence advantage) of a phenomenological model is that classification of seizure types through its parameters is based solely on the dynamic properties of the system, rather than assumptions on the underlying biophysical mechanisms as per other more physiological models such as the Jansen-Rit model (Houssaini et al., 2020). This is powerful, as epilepsy is believed to result from an instability of grey matter activity, observed consistently in humans and animals, even though the underlying pathological mechanisms often differ and remain unclear (Saggio & Jirsa, 2024). Studies have shown the ability of the Epileptor model to replicate not only many temporal features of seizure activity (V. Jirsa et al., 2023; H. E. Wang et al., 2023) but also spatial features of seizure activity, in that we can understand the spread of seizures in the brain (Moosavi et al., 2022). Individualised models can be created and then used to identify seizure-causing regions of the brain and predict the outcome of different surgical interventions (V. Jirsa et al., 2023; H. E. Wang et al., 2023).

Source localisation during epileptic seizures (ictal source localisation) has long been of interest due to its potential to non-invasively identify seizure-generating regions and to characterise the spatiotemporal recruitment of cortical networks during ictal events. Unlike interictal epileptic discharges, ictal activity evolves dynamically over time, spreads beyond the seizure onset zone, and often exhibits lower signal-to-noise ratios due to patient movement (van Mierlo et al., 2020). Despite these difficulties, ictal source localisation offers important clinical advantages. The ultimate goal of epilepsy surgery is to eliminate the origin of seizures rather than interictal discharges, and these two phenomena do not necessarily have the same origin (Bartolomei et al., 2016). By providing a more objective interpretation of scalp EEG than visual inspection alone, ictal source imaging can yield valuable localisation information to guide surgical resection or the placement of intracranial EEG electrodes (van Mierlo et al., 2020). Consequently, accurate localisation of ictal sources is of high clinical relevance and has been increasingly investigated as a potential electrophysiological biomarker for the epileptogenic zone (Jiang et al., 2025), providing a complementary perspective to interictal source imaging and other components of the epilepsy diagnostic pipeline.

This paper presents a framework for generating cortical simulations of ictal spread and their corresponding EEG/MEG signals. The key rationale is the need for an objective, repeatable methodology to determine the accuracy of source localisation approaches, with a focus on the minimum-norm family that dominates contemporary clinical EEG/MEG practice (van Mierlo et al., 2020). These simulations aim to answer the following questions:

1. How do different modelling parameters and decisions affect what we see at the source (neural) level as well as the macroscopic EEG/MEG level?
2. Are these simulations able to serve as a useful source of truth to benchmark source imaging algorithms?
3. How well can existing source localisation methods recreate pathological brain activity?

## 2 Methods

### 2.1 Epileptor Model

This study used a common implementation of the Epileptor model, as implemented in the Virtual Brain Project (TVB) (Sanz-Leon et al., 2015), where the dynamical behaviour of epileptic seizures is characterised using five state variables (*x*_1_, *y*_1_, *x*_2_, *y*_2_, *z*), a dummy variable (*u*) within region *i*, and an input variable *J*_*i*_ representing the excitatory input into region *i* from all other regions (V. K. Jirsa et al., 2017). The Epileptor equations are

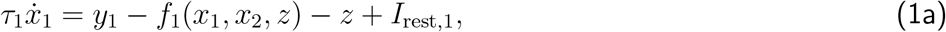

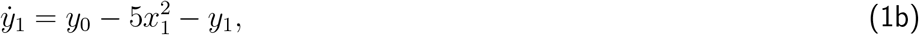

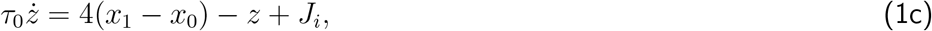

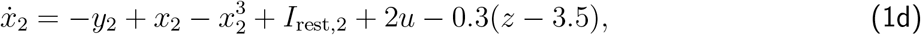

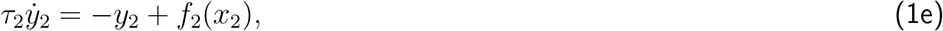

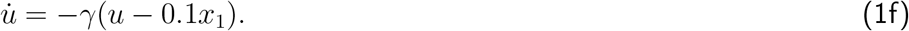

where

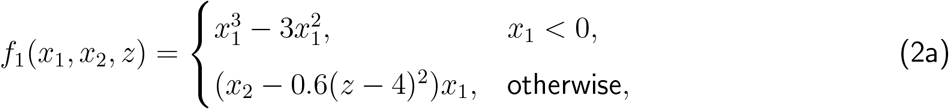

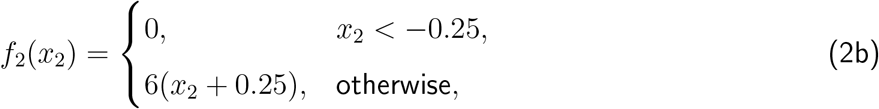

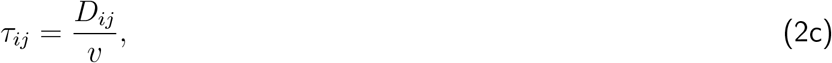

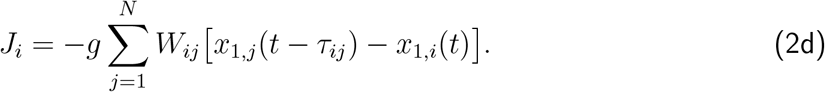

#### Representation as Dipoles

The Epileptor model generates time series representing dimensionless neural activity within a single region of the brain. To represent extracellular dipoles (*x*), the Epileptor equations were modified to

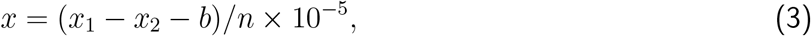

where *x*_1_ − *x*_2_ was used instead of *x*_2_ − *x*_1_, as by V. K. Jirsa et al. (2014), to ensure that the seizure state has a positive orientation, *n* is the number of dipole sources in the source space, and 10^−5^ converts to the physical units of A · m, consistent with the standard SI representation of neural current sources. In this formulation, *b* is a baseline constant, subtracted from the Epileptor model, to ensure that dipole sources maintain approximately zero baseline activity, as expected from the Debye shielding of ions in the electrolyte solution of the extracellular space (Nunez & Srinivasan, 2006).

#### Coupled Neural Mass Models for Seizure Spread

To generate seizure activity across the brain, this study used simulations of coupled masses of Epileptor models, as defined in Equation 1, through the coupling current *J*_*i*_. In this framework, each neural mass model corresponded to one cortical or subcortical region according to the Yan 1000 cortical parcellation atlas (Yan et al., 2023). The connectivity weight matrix (*W*_*ij*_ in Eq. 2d) was therefore defined by the structural connectome of an exemplar human subject (see Sec. 2.2). The axonal time delays *τ*_*ij*_ were defined by the 1019 region connectome streamline lengths (*D*_*ij*_) divided by a conduction speed of 3 m/s (Sanz Leon et al., 2013).

**Table 1.** Table of Variables and Parameters of the Epileptor.

| State variables |  | Interpretation |
| --- | --- | --- |
| $x_1(t), y_1(t)$ | | Fast timescale rapid discharges |
| $x_2(t), y_2(t)$ | | Intermediate timescale spike-and-waves |
| $z(t)$ | | Slow timescale transition between seizure and non-seizure |
| Other variables |  |  |
| $u(t)$ | | Dummy variable for low-pass filtering from $x_1$ to $x_2$ |
| $J_i(t)$ | | Coupling term describing total input into region $i$ from all other regions |
| Adjusted parameters |  |  |
| $x_0$ | $-2.1 \rightarrow -1.6$ | Node excitability |
| $g$ | $1.5 \rightarrow 2.0$ | Global coupling strength |
| $W_{ij}$ | $0.0 \rightarrow 1$ | Coupling weights defined by SCM into region $i$ from $j$ |
| $D_{ij}$ | $0.0 \rightarrow 248.5$ | Streamline lengths between regions (mm) |
| $\tau_{ij}$ | $0.8 \rightarrow 82.8$ | Time delays between regions (ms) |
| Fixed parameters |  |  |
| $I_{rest,1}$ | 3.1 | Constant injection current for $x_1$ |
| $I_{rest,2}$ | 0.45 | Constant injection current for $x_1$ |
| $y_0$ | 1 | Threshold constant for $y_1$ |
| $\gamma$ | 0.01 | Time constant of low-pass filter |
| $\tau_1$ | 1 | Time constant of fast activity |
| $\tau_2$ | 10 | Time constant of spike-and-waves |
| $r = 1/\tau_0$ | 0.00015 | Temporal scaling of long timescale activity |
| $v$ | 3 | Axon conduction velocity ($\text{ms}^{-1}$) |
| $\sigma^2$ | 0.0001 | Stochastic noise term (variance) affecting frequency of inter-ictal spikes and seizures |

A Heun’s stochastic integrator was used with Gaussian additive noise (∼ *N*(0, *σ*^2^)) added to the variables *x*_2_ and *y*_2_ (V. K. Jirsa et al., 2014; Moosavi et al., 2022). An integration step-size Δ*t* = 0.05 was used. To match the temporal properties of focal seizures, the timescale of the model output was scaled such that 256 time steps correspond to 1 s, as similar to V. K. Jirsa et al. (2017). Seizure simulations were set to have a 1 minute duration to allow sufficient ictogenic activity to develop. To generate each simulation, a single given region (i.e., represented by one Epileptor model) was first defined as the ‘epileptogenic zone’. All regions other than the epileptogenic zone were set to have an excitability of *x*_0_ = −2.1; i.e., tuned to be at the cusp of a bifurcation (Houssaini et al., 2020; Moosavi et al., 2022). To trigger a seizure, the excitability *x*_0_ for the specified epileptogenic zone was increased to −1.6. The global coupling strength was set at *g* = 1.5 or *g* = 2 to obtain two different degrees of seizure spread.

In total, 2038 simulations were generated to produce a dataset with variations in seizure morphology by changing the specified epileptogenic zone and the global coupling strength. The carpet plots of Fig. 1a and Fig. S4 depict the seizure propagating across the simulated brain regions. With higher coupling strength (*g* = 2), all simulations produced seizures that spread throughout all brain regions (as in Fig. 1). Intermediate coupling (*g* = 1.5) produced seizures that only partially spread through the brain in the 1 minute seizure duration (as in Fig. S4). Partial seizure spread occurred as a result of three effects of lower connectivity strength: i) reduced gain in external current between regions, ii) slower region recruitment, and iii) the refractory period of each region (preventing too many regions from undergoing a seizure together). The result is a lower net current between regions, meaning that some regions never receive enough current to drive a transition into a seizure state.

**Figure 1.**
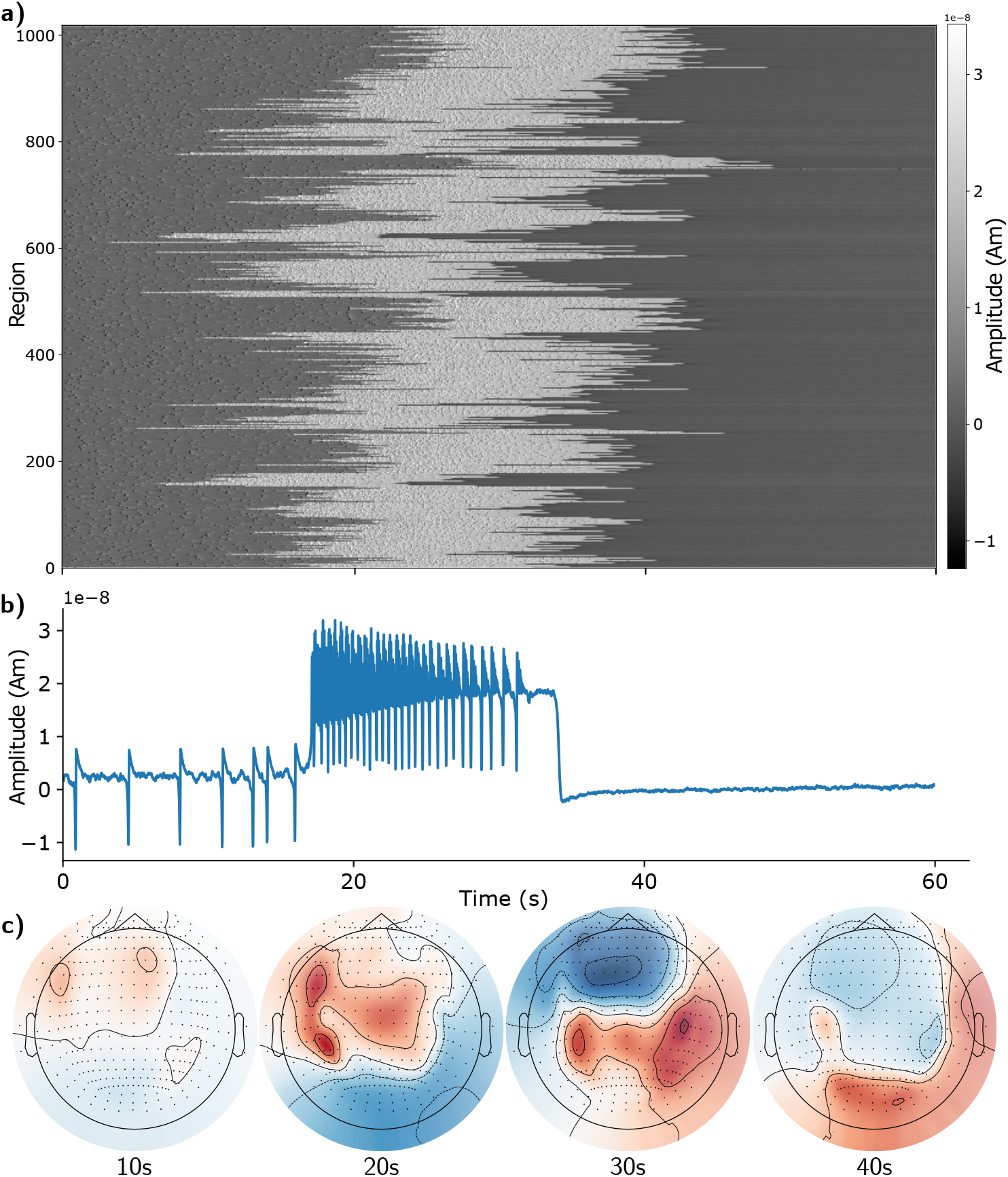
Epileptor model seizure originating from the left temporal pole. a) Carpet plot (Siu et al., 2022) of a simulated seizure originating in the left temporal pole with coupling strength *g* = 2. White regions indicate high-amplitude epileptiform activity. At this coupling level, seizure dynamics propagate across the entire brain network. b) Representative Epileptor time series showing key seizure features: interictal spikes (∼ −10 nAm), spike–wave discharges (∼ 20 nAm) with a DC shift, and a postictal refractory period without interictal spikes. The Epileptor equations were adjusted (Eq. 3) so that baseline activity is centred near zero and seizure activity appears positive. c) Example EEG scalp topography obtained by projecting the source activity in panel a to sensor space using the forward model (Eq. 4).

### 2.2 Structural Brain Data

Structural brain data from the Human Connectome Project (HCP) (Van Essen et al., 2013) was used, with HCP Subject 100206 (HCP100206) chosen as the primary subject for this work. The T1 MRI image with the skull included was used to generate surface meshes in the subject’s native space for the skin, skull, and cortex using Freesurfer’s recon-all (Fischl, 2012). The individual subject’s cortical mesh was aligned to the fsaverage template mesh, which consisted of 163,842 vertices per hemisphere.

#### Parcellation Atlas

Each vertex was attributed to one of the cortical regions of a specified parcellation atlas. Three cortical parcellations were considered: the Yan 1000 region homotopic atlas (Yan et al., 2023), the Glasser 360 region atlas (Glasser & Van Essen, 2011), and the Desikan-Killiany 68 region atlas (Desikan et al., 2006). The Yan 1000 atlas was selected because its finer parcels provide a more direct correspondence between neural mass simulations and dipole source space, avoiding the overly large cortical patches produced by coarser parcellations. The location of each cortical region was defined as the centroid of its vertex coordinates. In addition to the 1000 cortical regions, 19 subcortical structures defined by the cifti grayordinate template (Glasser et al., 2013) were also included, resulting in a 1019-region parcellation of the entire brain.

#### Structural Connectome

The individual’s structural connectome was computed via a diffusion MRI tractography pipeline detailed in previous work (Mansour L et al., 2021, 2023) using the MRtrix3 software (J.-D. Tournier et al., 2019). The fibre orientation distributions (FODs) were computed from tissue-specific response functions via Multi-Shell Multi-Tissue Constrained Spherical Deconvolution (Dhollander et al., 2016; Jeurissen et al., 2014). A total of 5 million tractography streamlines were estimated by an anatomically constrained probabilistic tractography algorithm based on second-order integration over FODs (Smith et al., 2012; J. D. Tournier et al., 2010). Notably, streamlines were randomly seeded from the gray matter white matter interface. A radial search with a maximum radius threshold of 4 mm was used to map streamlines to the regional parcellation, resulting in a 1019×1019 connectivity matrix of streamline counts. The matrix of streamline counts was then normalised by the maximum streamline count to end in a normalised connectivity matrix with entries between 0 and 1. In addition, a 1019×1019 streamline length matrix was computed to quantify average streamline lengths between all pairs of connected regions. To obtain the group-averaged connectivity matrix of streamline counts, the sum of streamlines between regions was taken for every patient, and the aggregate matrix is normalised between 0 and 1.

#### Dipole Source Space

The dipole source space is a representation of all dipole neural sources, located on the cortical surface, which can be summed to generate EEG/MEG data. These dipoles are nominally oriented normal to the cortical surface, but some flexibility can be allowed for the orientation to be ‘loose’ and deviate slightly from the normal. In this work, a discrete source space was used where the centre of each cortical region of the Yan 1000 parcellation was approximated as a dipole. Seizure (Epileptor neural mass model) simulations were mapped directly (1-to-1) to the source space based on the cortical parcellation of the subject.

### 2.3 Mapping to Sensor Space with MNE Python

Having generated simulated activity at the mesoscopic source level of brain regions, these simulated sources were then mapped to their corresponding signals in EEG/MEG sensor space using using Open-MEEG’s (Gramfort et al., 2010) implementation of the Boundary Element Method (BEM) for the forward problem with the corresponding forward model for the subject. Under the quasi-static assumption (Hallez et al., 2007; He et al., 2018), this transformation can be described by the linear model

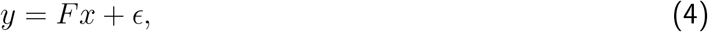

where the transformation matrix *F* ∈ ℝ^*M ×N*^ (known as the forward matrix or lead field matrix) maps any neural activity in source space *x* ∈ ℝ^*N ×T*^ to sensor space *y* ∈ ℝ^*M ×T*^. The dimensions represent the number of sensors (*M*), the number of possible sources in the brain (*N*), and the number of time steps (*T*), while *ϵ* is a matrix that represents the overall measurement noise for each time step. For the purposes of EEG source localisation, the quasi-static condition holds true, as it has been shown that charge does not accumulate in the extracellular space for the frequency range of signals measured with EEG Hallez et al., 2007, meaning that, for each instance in time, all electric and magnetic fields are dependent only on the active electric source Nunez and Srinivasan, 2006. The BEM model consisted of three concentric layers, with standard conductivity values assigned as: 0.3 S*/*m (skin), 0.006 S*/*m (skull), and 0.3 S*/*m (cortex) (Gramfort et al., 2014). The number of sensors varies depending on the EEG-spacing montage used, such that 10-20 spacing involves 21 electrodes, 10-10 spacing involves 88 electrodes, 10-05 spacing involves 339 electrodes. A visualisation of the sensors, dipole sources, and head surfaces can be found in Fig. S8.

To simulate the challenge of measurement noise contamination, Gaussian white noise was added to the EEG signal with signal-to-noise-ratios (*ρ*) of 3 dB, 10 dB, or None (signifying that no white noise was added). The standard deviation of the noise is given by

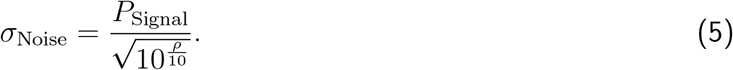

During the source inversion stage, the regularisation parameter (*α* or *λ*^2^) was set to 10^−*ρ/*10^, based on a heuristic.

### 2.4 Performance Metrics

To assess the accuracy of source localisation within simulations, the following evaluation metrics were used.

#### Cosine Score *S*_*C*_

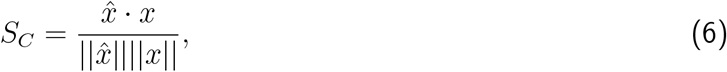

where *x* is the true signal and 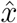 is the source estimate. Dependent on the figure, *S*_*C*_ was presented for each time point or averaged across time. The cosine score captures relative distributional accuracy, reflecting how well the relative pattern of activation across the source space matches the true activity.

#### Normalised Residual Sum of Squares *nRSS*

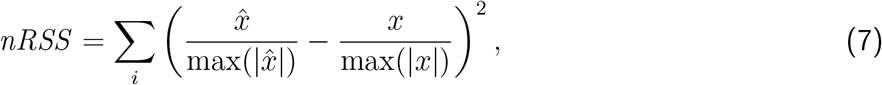

where the data was normalised using the maximal value, to compare across methods with differing units

#### Mean Squared Error *MSE*

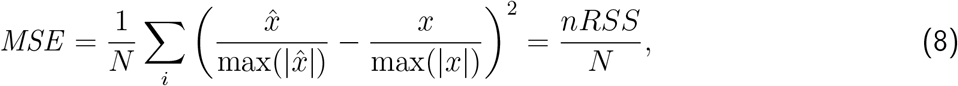

where *N* = 1000 is the number of dipole sources

#### Variance Explained *R*^2^

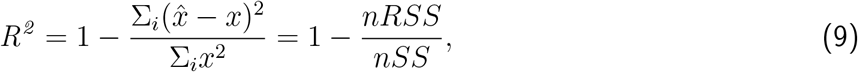

where *nRSS* is the normalised residual sum of squares and *nSS* is the normalised sum of squares over all possible sources *i* in source space. Note that for the simulated signals used here, a baseline constant was already identified and subtracted (see Eq. 3) such that the baseline for *x* is at 0

#### Region Localisation Error *RLE*

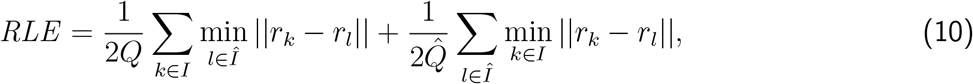

where *I* and 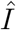 represent the true and estimated indices of active sources, respectively, *Q* and 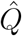 represent the number of true and estimated active sources, and *r*_*k*_ denotes the position of the *k*-th dipole source in space as per MNE-Python’s implementation (Gramfort et al., 2014). Sufficiently active sources were algorithmically identified to be those beyond Otsu’s Threshold (Otsu, 1979) as suggested by Sun et al. (2022, 2024). The region localisation error was computed across time, as well as at 50% of maximum EEG power, representing a sensible extent of seizure spread and the rising phase of a seizure (Plummer et al., 2019). Region localisation error captures spatial accuracy, reflecting how precisely the estimated source activity is positioned in space.

### 2.5 Source Localisation Algorithms

Having created plausible simulations of seizure spread at both the sensor and source levels, the simulations were used to evaluate the performance of four current source localisation approaches. These algorithms were implemented in MNE-Python and fall under the MN-family of source localisation algorithms, consisting of MNE (Hämäläinen & Ilmoniemi, 1994) (classical minimum L2 norm estimation), dSPM (Dale et al., 2000) (MNE with posthoc noise normalisation), sLORETA (Pascual-Marqui et al., 2002) (MNE with source-covariance normalisation), and eLORETA (Pascual-Marqui, 2007) (MNE with minimum localisation error weighting). All methods used an identical forward model, source space, depth, orientation, and regularisation parameters such that differences in performance reflected the inverse solution alone.

## 3 Results

In total, 2038 simulations were generated to produce a dataset with variations in seizure morphology by changing the specified epileptogenic zone (1019 possible regions) and the global coupling strength (two possible values). Fig. 2 provides a visual representation of the abilities of the MN-family to recreate the propagation of an example seizure simulation originating from the left temporal pole with coupling strength *g* = 2. The following sections outline insights into the performance of the four source localisation algorithms compared.

**Figure 2.**
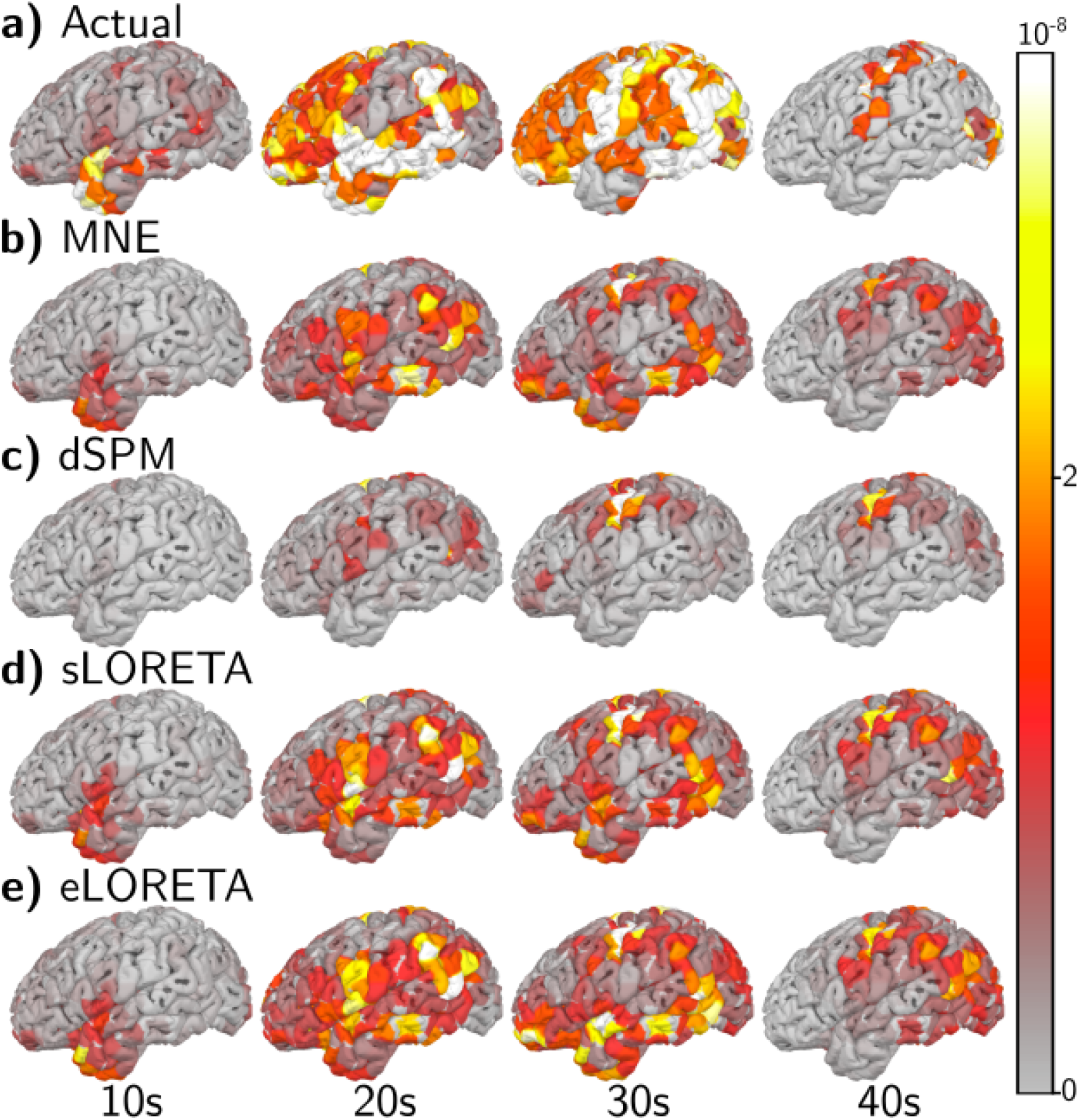
MN-family estimates for seizure originating from left temporal pole plotted on left hemisphere. The colour scale represents source amplitude, with white indicating the maximum amplitude and transitioning through yellow, orange, and red to transparent at zero. a) Ground-truth source activity generated using the Epileptor model. As the simulation progresses, seizure activity propagates from anterior to posterior brain regions. During this period, different cortical areas become active with varying spatial extents and patterns. b)-e) Algorithms from the MN-family predict more focal sources with regularisation parameter set according to the 10^−ρ*/*10^ heuristic. For visualisation and to improve performance, the orientation of the dipole was allowed to be loose for the MN-family, which we demonstrate to be near equivalent to taking the absolute value of a fixed orientation estimate.

### 3.1 Existing Approaches are Challenged by Irregular Sources

For a more direct comparison with previous literature, which usually focuses on one or a few sources of various extents (Cioppa et al., 2023; Giri et al., 2022; Hecker et al., 2021; Morik et al., 2024; Petrov, 2012; Rong et al., 2025), we computed a simple set of single-source simulations. These sources were generated with varying sizes such that, for each simulation, one cortical region of the 1000-region parcellation was selected as the centre. Regions within a certain radius were then selected as part of the entire patch. This simulation would also be more similar to the case of averaged interictal spikes.

Fig. 3 shows the performance metrics for existing, ‘off-the-shelf’ source localisation methods. The region localisation error of approximately 0.5–2 cm (Fig. 3a) was consistent with previous findings (Hauk et al., 2022; Hecker et al., 2021; Sohrabpour et al., 2020; Sun et al., 2022). Interestingly, the reduction in region localisation error and the increase in mean squared error indicate that reconstruction accuracy declines as the spatial extent of the source increases. This behaviour is likely driven by the way extended patches were generated; as patch size grows, the resulting sources become less circular and develop increasingly irregular boundaries (see Fig. S7). The minimum-energy constraints inherent to MN-family algorithms are biased against irregular shapes and instead favour compact, smoothly distributed sources. Furthermore, larger patch sizes mean that sources encompass more areas of cortical curvature, producing cancellation when dipole normals are oriented in opposite directions. We also observe point-spread artefacts (i.e., spatial “ringing”) and smoothing effects in Fig. S7, which are well-known and typically undesirable features of MN-family inverse solutions (Hauk et al., 2022).

**Figure 3.**
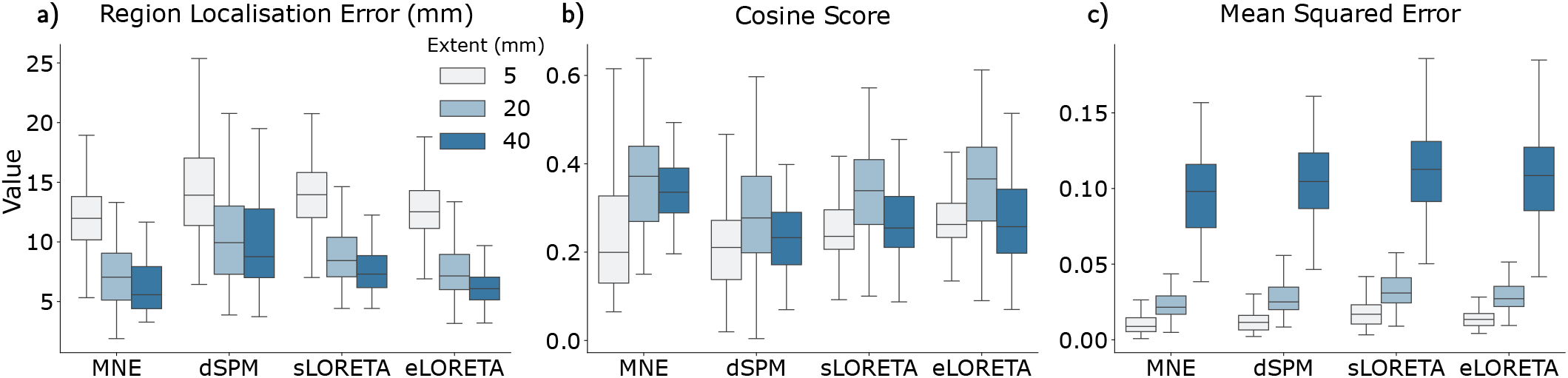
Single patch source localisation metrics. Source localisation results for varying extents were computed for off-the-shelf algorithms from MNE-Python. A 10-10 montage with 88 EEG electrodes was used. The box range shows the interquartile range and the median. Each subfigure graphs a different metric: a) region localisation error, b) cosine score, c) mean squared error. Each hue represents underlying sources of different extents (radius of 5mm, 20mm, 40mm).

### 3.2 Increasing the Number of Electrodes Allows Polarity to be Resolved

Fig. 4 shows the source localisation metrics across the simulated dataset, where the variable parameters were signal-to-noise ratio, number of electrodes, and algorithms. The previous section discussed the challenge of irregular and widespread sources for existing algorithms. This challenge is further evidenced by Fig. 4d, columns 2 and 3 (representing the EEG 10-10 and 10-20 montages, respectively). At 21 and 88 electrodes, the MN-family of methods was unable to accurately determine the polarity of sources, as evidenced by the variances explained being close to 0 and the relatively low cosine scores. At 343 electrodes, the MN-family was able to resolve the polarity of sources, leading to substantial increases in performance/accuracy. The estimation of polarity is discussed in more detail in Results Sec. 3.4 and Discussion Sec. 4.

**Figure 4.**
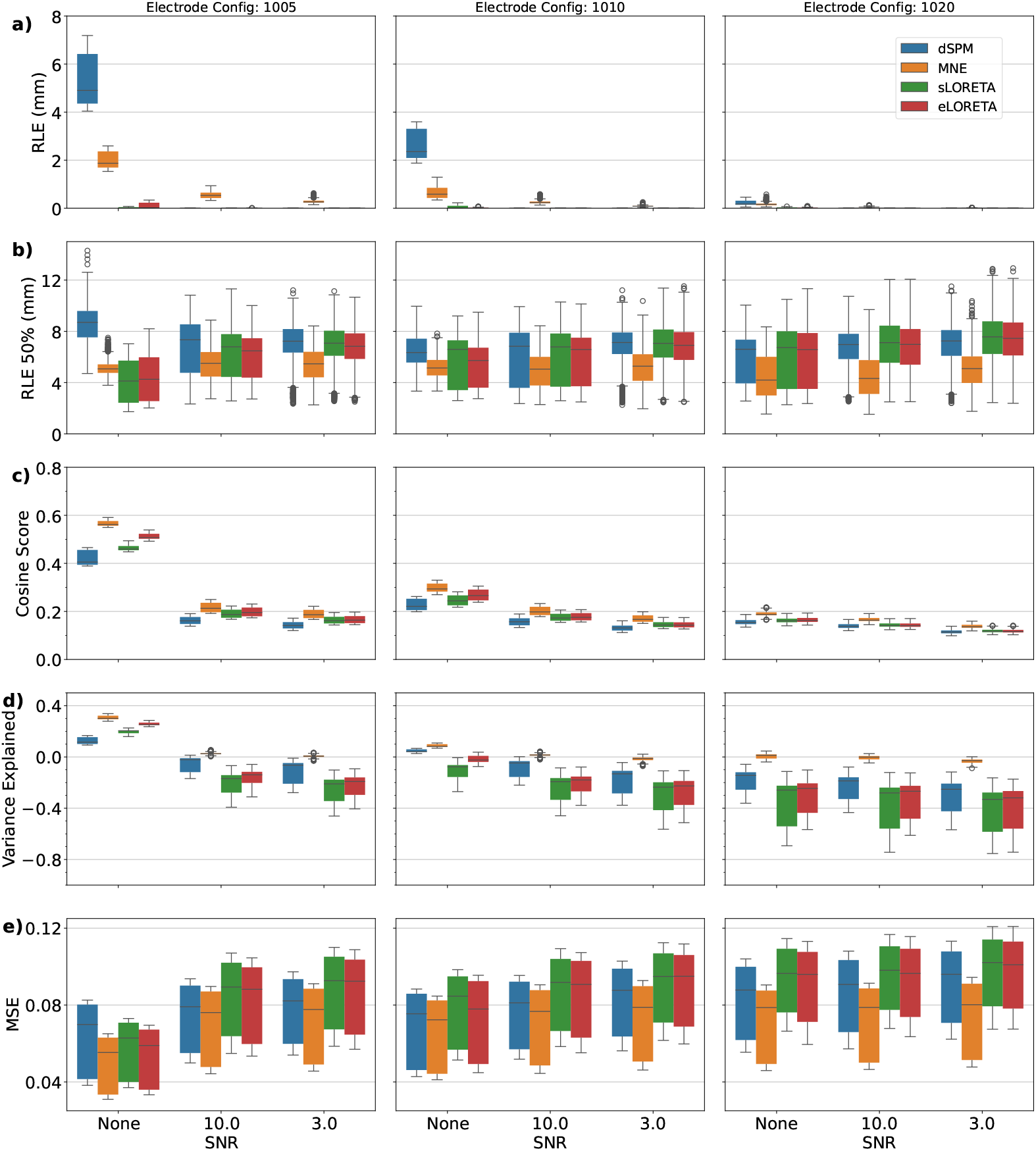
MN-family source localisation metrics. Each algorithm from the MN-family is represented by a different colour: dSPM (blue), MNE (orange), sLORETA (green), and eLORETA (red). Each row, labelled a–e, displays a different metric: region localisation error, region localisation error at 50% of EEG max power, cosine score, variance explained, and mean squared error. Each column corresponds to a different electrode configuration: ‘10-05’, ‘10-10’, and ‘10-20’ spacing. Within each sub-figure, the x-ticks represent the signal-to-noise ratio: none (no noise), 10 dB, and 3 dB. Metrics are computed with a ‘fixed’ orientation, meaning that dipoles are assumed to be oriented perpendicular to the cortical surface.

Considering the RLE at 50% of EEG power (Fig. 4b), performance appeared stable across electrode configurations at approximately 6 mm on average. This RLE signifies the reasonable spatial accuracy of the source localisation algorithms, and is in line with previous findings of (Hecker et al., 2021; Pascarella et al., 2023; Sharma et al., 2019). The RLE showed that the best results were obtained with no noise, and SNRs of 10 and 3 showed minimal differences after regularisation. dSPM performed poorly relative to the other algorithms for this metric, except for electrode configuration 10-20.

In terms of cosine score (Fig. 4c) and variance explained (Fig. 4d), both follow similar qualitative trends given that they are closely related metrics. The distinction is that variance explained has a wider dynamic range and is sensitive to both magnitude and alignment, whereas cosine score is bounded and, therefore, less sensitive to variation at higher (∼1) and lower (∼0) performance levels, and does not consider magnitude. Under noisy conditions or montages with 88 or fewer electrodes, the metrics indicate poor localisation results.

Overall, the mean squared error (Fig. 4e) was shown to increase by approximately 30% from no noise to 10 dB noise, with only a slight further increase from 10 dB to 3 dB for the 10-05 montage. These jumps in error as noise increased were less pronounced for simulations with fewer electrodes, likely because the baseline errors (in noise-free conditions) were already higher. This pattern, where the error increases quickly and then plateaus, likely signifies a saturation in error.

### 3.3 Localisation Error Varies Throughout the Seizure

Fig. 5 illustrates that region localisation error varies throughout a seizure, following a characteristic temporal profile. Immediately before and during the initial recruitment of seizure activity (0–10 s), the RLE is relatively high with substantial variability, indicating reduced confidence and stability in the source localisation estimate during the initial distributed activation. As the seizure evolves into the mid phase (defined as when 50% of maximum EEG power is reached (Plummer et al., 2019), typically occurring at around 10–20 s into the simulations), the localisation error decreases markedly and remains at its lowest level during the sustained ictal period (around 20–40 s). This indicates that source localisation becomes more stable when the spike waveforms are better established and the signal-to-noise ratio is stronger. Toward the end of the seizure (¿50 s), the RLE increases again, consistent with the broader spatial dispersion and complex propagation patterns in late ictal dynamics. Overall, these results suggest that the mid-ictal stage provides the most reliable signal for source localisation, whereas both early recruitment and late spread reduce localisation accuracy due to rapidly changing and irregular source configurations.

**Figure 5.**
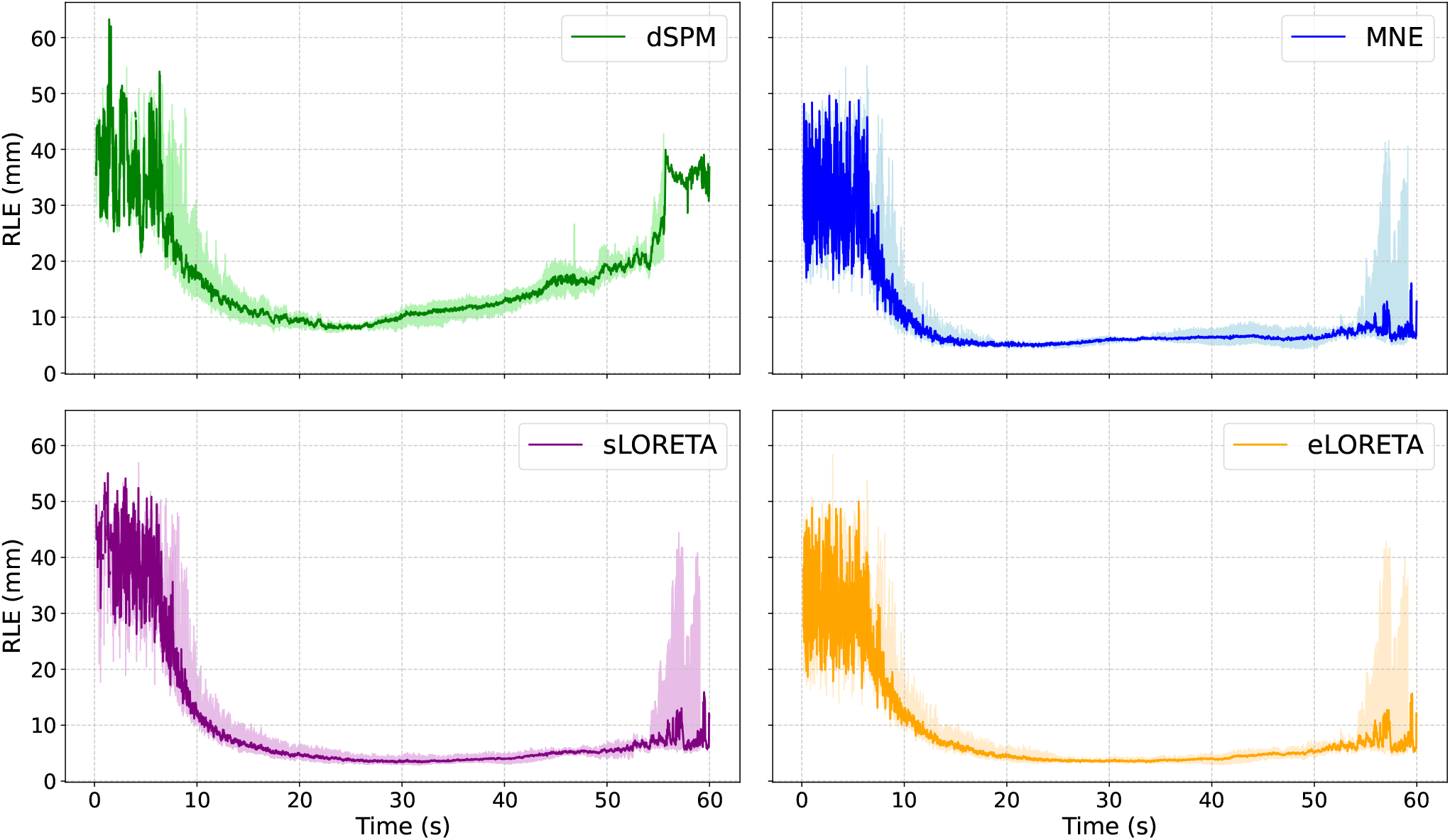
Region Localisation Error Over Time for algorithms in the MN-Family. The solid line shows the median region localisation error and the shaded region indicates the interquartile range across simulations. The mid-phase of a seizure is defined to start when 50% of maximum EEG power is reached, typically occuring at around 10-20 s into the seizure, depending on the simulation. Otsu’s threshold was used for calculating RLE shown, but the results remained consistent across thresholds from 10% to 60% for a SNR of 10.

### 3.4 Effect of Dipole Orientations

Allowing the orientation of the dipoles to be free to deviate from the normal of the cortical surface (also referred to as loose orientation) is a common approach in source localisation (Gramfort et al., 2014; Lin et al., 2006). In this work, the dipoles were oriented normal to the cortical surface for the simulation. However, as shown in Fig. 6 and Fig. S5, allowing a deviation from the normal of the cortical surface, by enabling the dipole deviation to be greater than zero, increased performance for all source localisation methods. The standard explanation would be that the loose orientation provides extra degrees of freedom to account for small errors with the forward model, mesh discretisation, downsampling, etc. To test if this is the case, we consider the effects of different variations of implementing loose vs. fixed orientation using the eLORETA algorithm, as shown in Fig. 6. Sequentially, the shades of blue represent fixed orientation, and the warm colours represent settings of loose orientation.

**Figure 6.**
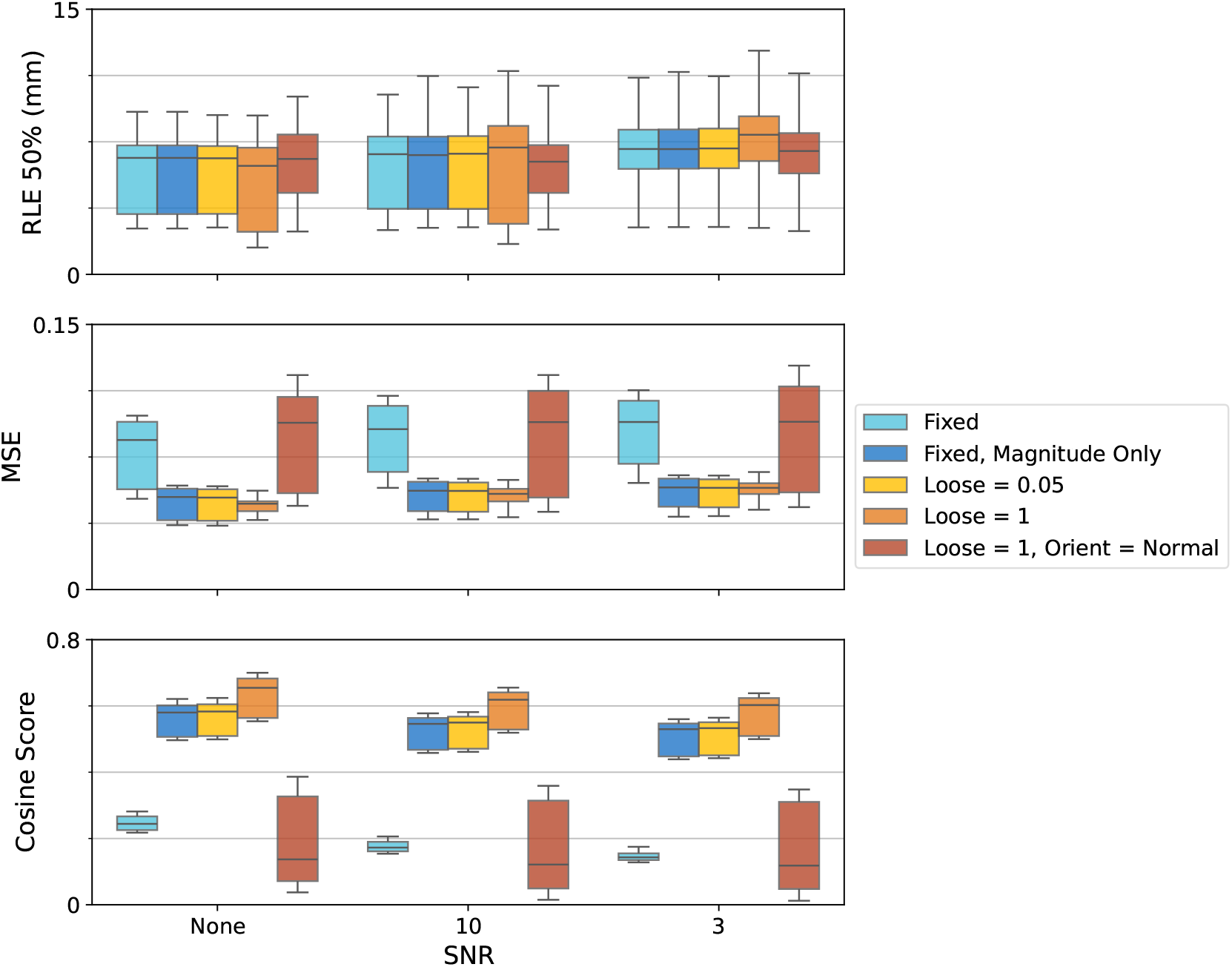
Results of eLORETA when adjusting the loose orientation parameter. Each row is a different metric (region localisation error at 50% EEG power, mean squared error, cosine score), and the x-axis indicates the signal-to-noise ratio in dB. Metrics are computed with a ‘loose’ orientation, which controls the degree to which dipole orientations are allowed to deviate from the normal of the cortical surface. Different loose orientation parameter settings are represented by different colours: Light blue = Fixed orientation, Dark blue = Fixed orientation with taking the absolute value, Gold = Minimally loose orientation (parameter set to 0.05), Orange = Loose orientation, Mahogany = Loose orientation, but taking only the normal to the surface component.

In Fig. 6, comparing the fixed orientation (light blue) with loose orientation (gold) demonstrates how setting a loose orientation significantly improves source localisation performance, with mean squared error reducing from 0.077 to 0.048 under no noise conditions. However, the gold boxplot demonstrates that the performance increase can also be seen by simply taking the absolute value of the fixed orientation estimate. Further increase of the loose parameter (comparing gold to orange) appears to only slightly increase the performance for eLORETA, with mean squared error improving from 0.052 (gold: loose=0.05) to 0.048 (orange: loose=1) under no noise conditions. Essentially, what this means is that the removal of the direction (and only taking the magnitude) produces the bulk of the performance increase, rather than the allowance of deviation to the cortical surface. Indeed, the mahogany boxplot displays the estimate and metric when only considering the orientation component normal to the surface results in relatively poor performance. This is supported by region localisation error, a polarity agnostic measurement, remaining relatively stable across changes to the loose parameter. This is not necessarily a flaw with loose orientation, but an artefact of vector estimation. After finding the loose source estimate, there is a need to compare it with the true source, which is oriented normal to the cortical surface. The typical method used to compare the estimated with true sources, which is the default in MNE-Python, is to condense the vector into a magnitude and compare metrics, and this is what was performed to generate the results for the loose orientations in Fig. 6 (warm colours).

## 4 Discussion

This study introduced a benchmarking framework for source localisation algorithms using simulated seizures to evaluate performance against known onset regions. Simulations were able to capture key features of clinical epileptic seizures in both the source space and the resulting EEG sensor space, providing a realistic test bed for algorithmic evaluation and paving the way for future clinical applications. The results showed that existing source localisation approaches performed reasonably well under idealised conditions (343 electrodes, no measurement noise), but their accuracy degraded substantially under more realistic scenarios involving noise and lower-density montages. The primary contributor to this performance decline was a reduced ability to recover the correct polarity of the underlying sources, even when the spatial localisation itself remained relatively accurate. For the purposes of localising the epileptogenic zone or tracking regional recruitment during seizures, spatial accuracy is typically the dominant consideration. Nonetheless, improving polarity reconstruction remains an important area for future development, particularly for studies concerned with mechanistic insights of seizure dynamics or more generalised hierarchical relations in the brain, as discussed later.

### Resolving the Polarity of Dipole Sources

The current results showed that the MN-family of source localisation algorithms was unable to reliably reconstruct the correct polarity of dipole sources under typical EEG conditions, particularly when using lower-density montages. In EEG and MEG source localisation, polarity reflects the orientation of the underlying current dipole relative to the sensors. Although forward models preserve this orientation, inverse solutions often fail to recover it due to the ambiguity of multiple source configurations explaining identical sensor patterns, leading many pipelines to prioritise spatial accuracy over polarity (Gramfort et al., 2014). For applications such as identifying the epileptogenic zone (or, more broadly, any task focused on where activity originates), spatial precision is indeed far more critical than recovering dipole orientation, especially given susceptibility of polarity to noise, head-model inaccuracies, and orientation ambiguities, as demonstrated in the preceding simulations.

Nevertheless, polarity can carry meaningful physiological information. The polarity represents the direction of current flow across the cortical surface and through the cortical laminae, with implications on the hierarchical (such as feedforward or feedback) relationships between areas in the brain (Ahlfors & Wreh, 2015; Ahlfors et al., 2015; Lankinen et al., 2024). Polarity may also be essential when interpreting functional connectivity (Coelli et al., 2023; Haufe & Ewald, 2019), phase relationships (Haufe et al., 2011; Koller et al., 2024), or propagation direction (Koller et al., 2024), all of which are vital components of understanding the mechanisms underlying seizures or other generalised wave-like brain activity. Therefore, incorrect signs can invert inferred interactions or misrepresent the temporal ordering of neural activations. As such, the importance of resolving the correct polarity of the dipole sources should be guided by the scientific or clinical question being asked.

### Optimal Timing for Epileptogenic Zone Localisation

This study identifies the mid-phase of a seizure, the period following 50% mean global field power, as the optimal time point for EZ localisation (Fig 5). This aligns with established interictal practices (van Mierlo et al., 2017; G. Wang et al., 2011), yet appears to contrast with findings by Plummer et al. (2019), who favoured the early ictal period, as defined by the earliest time point at which 90% of the signal variance could be reconstructed. This discrepancy is primarily methodological. While the 90% variance criterion demands highly regularised, noise-robust inversion, our simulations show that stable localisation under more lenient regularisation emerges between ∼8 s (when the RLE became more stable at each time point) and ∼10 s (the “elbow” point at which the RLE dropped sharply and began to plateau). Both perspectives suggest that accuracy requires sufficiently organised seizure activity, however, this creates a fundamental clinical trade-off. By the time the signal reaches the stability required for reliable inverse reconstruction, the seizure has typically propagated to secondary regions. Consequently, the clearest reconstruction conditions may spatially misrepresent the underlying pathology by masking the true onset zone with widespread recruitment, reinforcing the necessity of capturing the earliest possible ictal generators.

### Current Methods of Evaluation

Current methods of evaluating source localisation results (as estimated from mesoscopic recordings) can be categorised into three groups:

1. Comparison with simultaneously measured sources,
2. Comparison with empirically inferred sources (i.e., using surgical resection outcomes to infer epileptic sources), and
3. Comparison with simulated sources.

A recent and promising development within the ‘simultaneous measurement’ category is the availability of datasets that combine simultaneous EEG or MEG with intracranial recordings, such as sEEG (Mikulan et al., 2020; Pascarella et al., 2023). These datasets offer a unique opportunity to compare non-invasive estimates with invasive ground truth. For example, Parmigiani et al. (2022) published an open sEEG dataset based on cortico-cortical evoked potentials from bipolar stimulation. However, to our knowledge, there are currently no published studies that evaluate source imaging approaches with simultaneous invasive and non-invasive measurements of neural activity. Key factors limiting the application of simultaneous recording include technical complexity, restricted spatial coverage with sEEG electrodes, and the presence of stimulation (used to generate clear sources) artefacts dominating both sEEG and EEG signals. Several studies have evaluated techniques using the stimulation artefact itself as the underlying ‘source’ (Mikulan et al., 2020; Pascarella et al., 2023; Unnwongse et al., 2023), but these are limited to evaluating localisation performance for a single source (the stimulation artefact), which differs from neural activity in terms of amplitude, waveform, and physical origins.

A second strategy involves evaluating the localisation of the epileptogenic zone in patients using surgical outcome as a proxy for ground truth (Plummer et al., 2019; Sun et al., 2022). This method provides a clinically relevant benchmark, but the use of surgical success as a binary outcome measure may bias results in favour of methods that produce larger estimated zones, thereby increasing false positives. Finally, the most common benchmarking approach by far has been to simulate the underlying sources directly and apply a forward transformation to obtain a data set with both the underlying sources and the resultant EEG signal (Barzegaran et al., 2019; Hauk et al., 2022). This is the approach undertaken by this work, where a clear distinction is the use of biologically-inspired neural mass models to generate the simulations as opposed to relying on a small number of spatially discrete sources (Cioppa et al., 2023; Giri et al., 2022; Hecker et al., 2021; Morik et al., 2024; Petrov, 2012; Rong et al., 2025). Furthermore, while Sun et al. (2022) have also employed neural mass model simulations, their source localisation algorithm was a deep learning model that was both trained and evaluated on the same simulated environment. No such circularity occurs here.

### Limitations and future work

While simulations provide valuable ground truth for assessing methodological performance, they inevitably simplify the complexity of real neural dynamics and are unlikely to capture the full range of noise and variability present in real-world data. Specifically, in this work, while the Epileptor model is widely used and is capable of reproducing many spatial and temporal features of epileptic dynamics at both microscale and macroscale levels (V. K. Jirsa et al., 2014; Moosavi et al., 2022; H. E. Wang et al., 2023), in this implementation, it employs only one of the twelve theoretically defined dynamical transitions associated with seizure initiation and termination (Saggio et al., 2020). Moreover, assumptions made during simulation design, such as source extent, timescales, and propagation dynamics, may introduce biases in favour of specific modelling approaches. While absolute accuracy in simulation typically overestimates performance in real data, relative performance trends, such as the comparative ranking of methods or the impact of specific parameter adjustments, often generalise (Sargent, 2013). Although simulations cannot substitute for clinical validation, in domains such as source localisation, where a definitive ground truth is currently not available, they serve as a critical intermediary, helping to guide expectations, identify limitations, and narrow the space of viable methods prior to clinical deployment. Future work should complement simulation-based evaluation with other forms of real data (such as surgical resection results of epilepsy patients) to verify findings in vivo.

A primary limitation of this study is its focus on a single candidate human model. While this approach is sufficient for validating the utility of the neural mass benchmarking framework, it does not account for inter-subject variability. The spatial-temporal dynamics of seizure propagation are highly dependent on the underlying structural connectome and individual anatomical geometry (Moosavi et al., 2022). Consequently, subject to a relevant hypothesis, future research can use multiple subject templates to investigate how individual differences in brain network topology and head models influence the propagation of pathological activity and the comparative performance of inverse algorithms on such signals. Furthermore, while this work only tested Gaussian measurement noise, clinical recordings are characterised by the presence of biological artefacts, such as myogenic and ocular activity. Expanding the framework to include multiple subjects and biological artefacts will allow for a more robust evaluation of how source localisation methods perform across broader clinical populations and recording conditions.

### Conclusion

This study established a benchmarking framework based on biologically inspired neural mass models to evaluate the performance of EEG source localisation algorithms. Using the Epileptor model to generate synthetic seizures, this work provides a realistic testbed that accounts for the complex spatiotemporal dynamics, such as seizure recruitment and propagation, characteristic of clinical epilepsy. The findings obtained through this simulation framework demonstrated that while MN-family algorithms were spatially accurate under idealised conditions, they began to fail under typical clinical environments such as increased measurement noise and lower-density montages. Ultimately, the adoption of neural mass models or other more biophysical simulations for benchmarking source imaging approaches would shift the focus from simple spatial precision to a more nuanced understanding of how source inversion algorithms can represent various dynamical states and regimes in the brain.

## Data and Code Availability

Structural connectivity data is available from the Human Connectome Project (HCP) (Van Essen et al., 2013). Simulation and analysis code for the evaluation pipeline can be found at https://github.com/Spokhim/sl_sim_framework.

## Author Contributions

P.H.S (Conceptualisation: Lead; Data curation: Equal; Formal analysis: Lead; Investigation: Lead; Methodology: Lead; Visualisation: Lead; Writing – original draft: Lead; Writing – review & editing: Equal), P.J.K. (Investigation: Supporting; Supervision: Equal; Writing – original draft: Supporting; Writing – review & editing: Equal), S.M.L. (Data curation: Equal; Writing – review & editing: Supporting), A.S. (Supervision: Equal; Writing – original draft: Supporting; Writing – review & editing: Equal), L.K. (Writing – original draft: Supporting; Writing – review & editing: Equal), M.J.C (Formal analysis: Supporting; Supervision: Equal; Writing – original draft: Supporting; Writing – review & editing: Equal), D.B.G. (Supervision: Equal; Writing – original draft: Supporting; Writing – review & editing: Equal)

## Acknowledgements

This research was supported by The University of Melbourne’s Research Computing Services, the Petascale Campus Initiative, and an Australian Government Research Training Program (RTP) Scholarship. This research was funded by the Australian Research Council (Discovery Project DP200102600).

## Supplementary Material

Supplementary material for this article is available with the online version.

## S1 Supplementary Material

### S1.1 Epileptor Model Mechanisms

Fig. S1a displays the three subsystems of the Epiletor model that each operate at a different timescale. The fast subsystem (*x*_1_, *y*_1_ - blue) is based on a modified Hindmarsh–Rose model (Hindmarsh & Rose, 1984) and is responsible for the fast oscillatory activity observed during seizures. This subsystem exhibits bistability between a stable fixed point (representing the interictal, resting state) and a stable limit cycle (oscillatory activity representing the ictal state).

The fast subsystem is coupled to a slow subsystem (*z* - green), which governs transitions into and out of the ictal state through a hysteresis loop under appropriate parameter regimes (Saggio & Jirsa, 2024). The phase diagram of Fig. S1b displays this hysteresis loop as a clockwise trajectory, forming the core mechanism of the Epileptor model.

Finally, the intermediate-timescale subsystem (*x*_2_, *y*_2_ - orange) operates near a saddle-node on an in-variant circle (SNIC) bifurcation (Houssaini et al., 2020). Each crossing of this bifurcation point by the slow subsystem generates a distinct spike through following an invariant circle trajectory. During the preictal period, these preictal spikes reflect increased excitability of the intermediate subsystem near seizure onset. During the ictal period, the intermediate variables modulate the fast variables such that together they produce the characteristic spike–wave complex (*x*_1_ − *x*_2_ in Fig. 1b).

### S1.2 Analysis of the EEG Activity Generated by the Epileptor Model

As with all simulations, due diligence must be performed to ensure that the simulations meet our requirements. Fig. S2 shows the EEG trace of the sample seizure originating from the thalamus. A neurologist reviewer confirmed that the simulated EEG exhibited clinically consistent features, including focal on-set, spatial spread, and temporal evolution with characteristic slowing. Fig. S3 visualises the spectral characteristics of the sample simulation. From Fig. S3a depicts the spectrogram for a single electrode, demonstrating how most power is concentrated at the very low frequencies, although some patches of power are also evident at higher frequencies. In terms of temporal information, it is clear when the seizure is occurring. In Fig. S3b, taking the log of the power reveals two major deviations from the brain’s natural 1/*f* ^*α*^ power scaling laws (Donoghue et al., 2020) in the low-frequency and the 15-20 Hz range. These peaks coincide with the ‘DC shift’ and the ‘fast oscillatory activity’ of population *x*_1_ in the Epileptor model.

---

**Figure S1.**
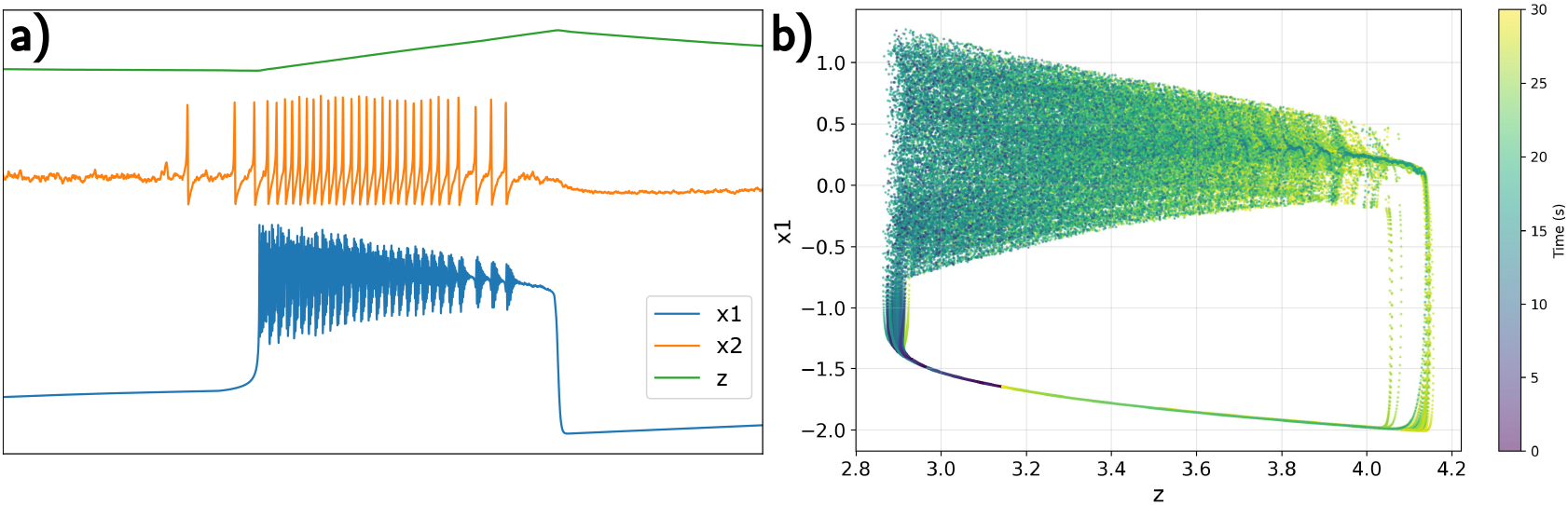
Mechanisms of the Epileptor model. a) The Epileptor model can be viewed as the interaction among three subsystems operating at different timescales: *x*_1_ (blue) at the fast timescale, *x*_2_ (orange) at the intermediate timescale, and *z* (green) at the slow timescale. b) *x*_1_ − *z* phase diagram where the colour represents the time of the trajectory. We see that, for a system initialised at (*x*_1_, *z*) = (−1.6, 3.1), it starts on a fixed point and then moves clockwise in the phase diagram (seen by the colour change from purple to dark green) to reach the unstable limit cycle on the upper portion of the loop. The system then continues along the hysteresis loop until it reaches the fixed point, with the cycle continuing if the simulation was continued.

**Figure S2.**
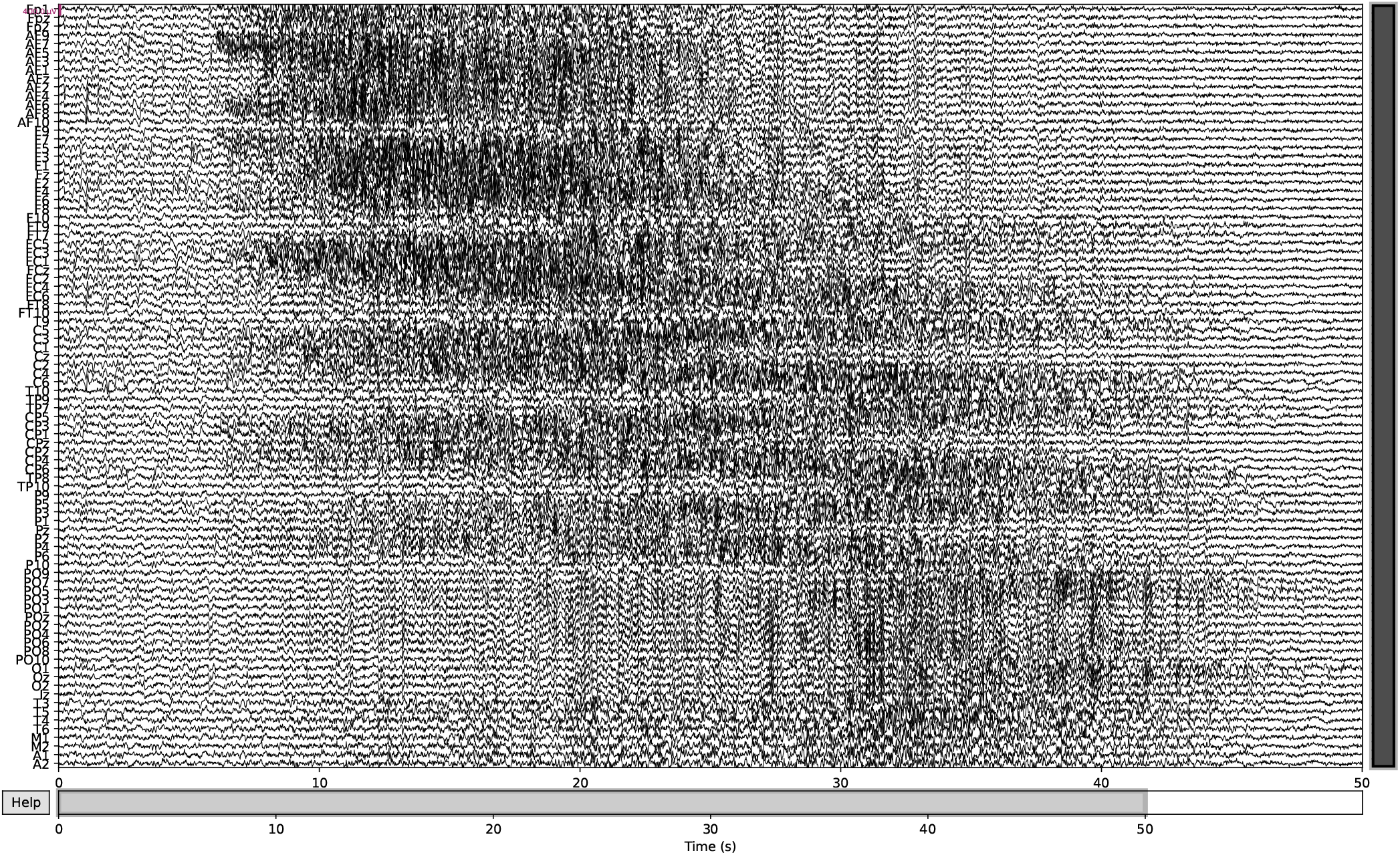
EEG trace of simulation with SNR of 10 dB. Each row represents the simulated EEG activity at each labelled electrode. Increase volatility can be seen in areas undergoing epileptic activity. The simulated EEG exhibited clinically consistent features, including focal onset, spatial spread, and temporal evolution with characteristic slowing.

**Figure S3.**
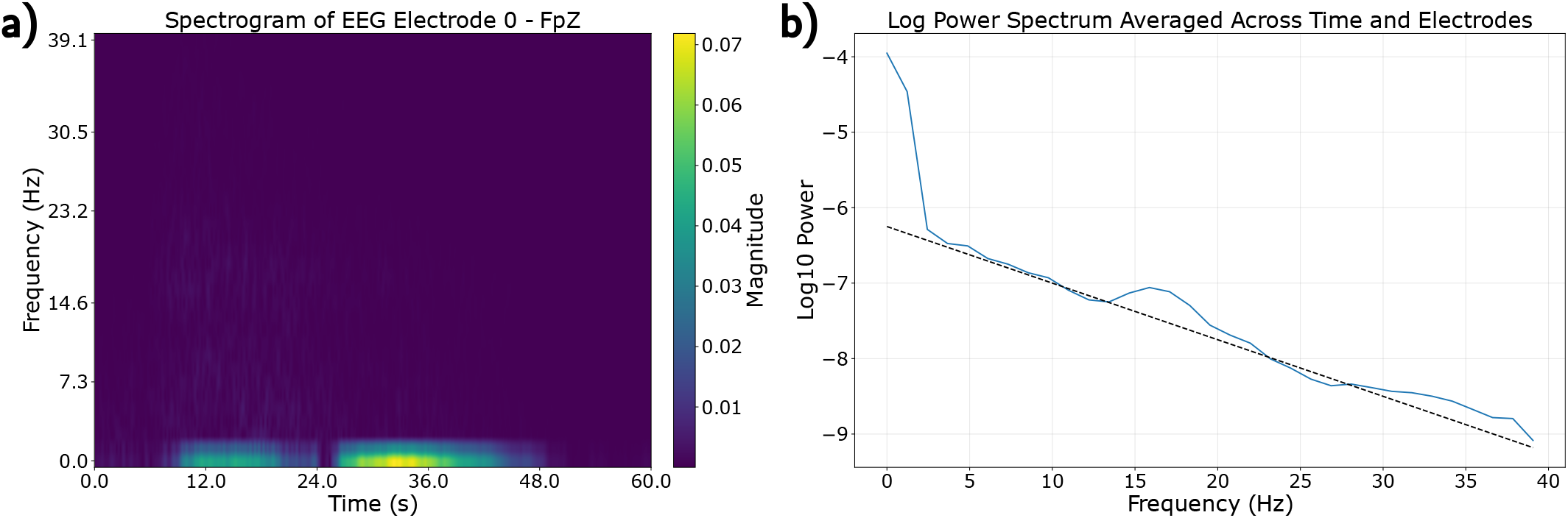
EEG simulation frequency spectrum. Spectral properties for the thalamic seizure simulation are graphed. a) Spectrogram for EEG electrode 0 - Fpz across time and frequency. b) Log power spectrum averaged across time and electrodes. The dotted line represents a 1*/f* ^*α*^ power scale (*α* = 3*/*40 in this instance).

**Figure S4.**
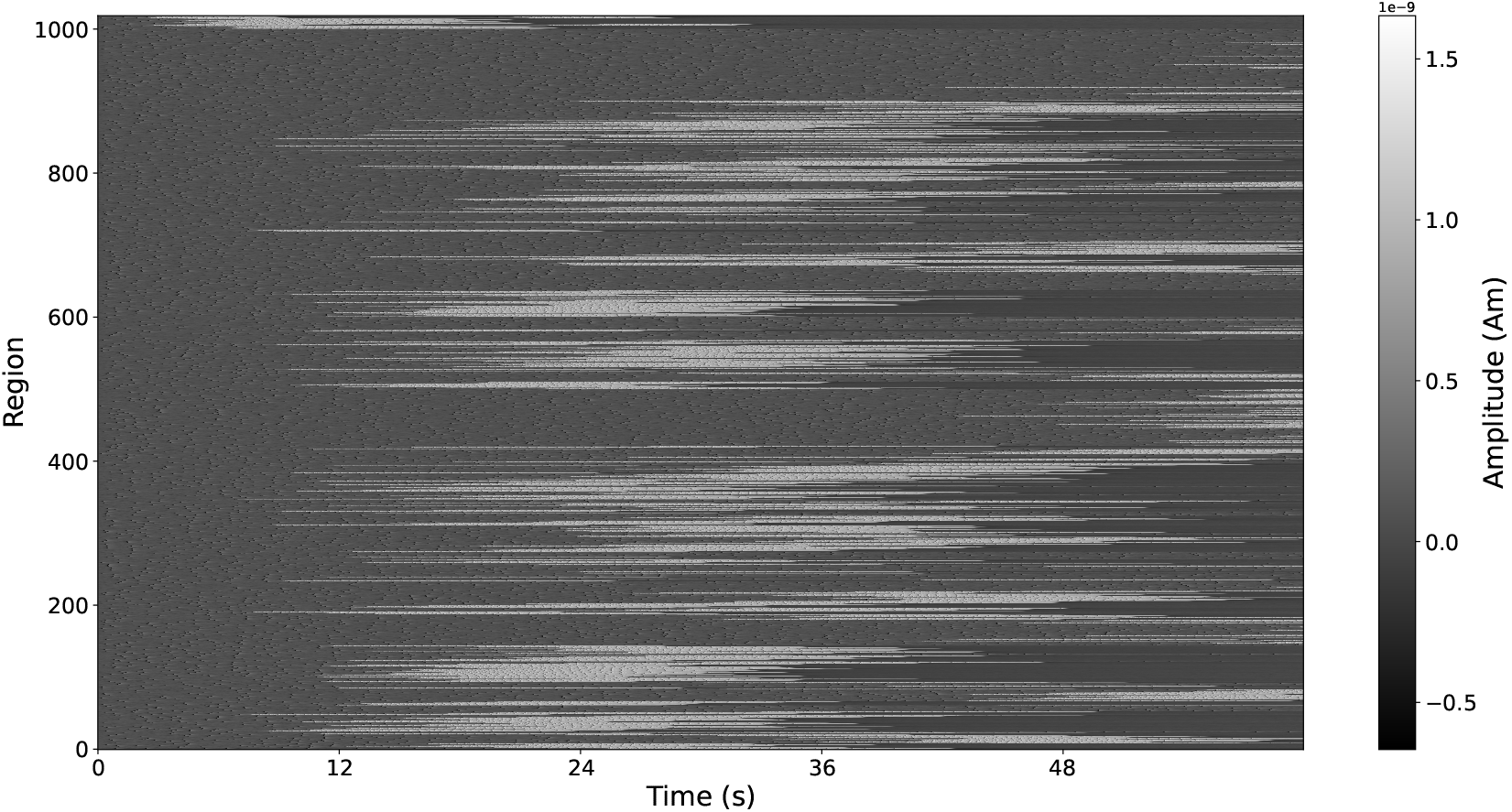
Carpet plot of seizure originating from thalamus with partial spread. Partial spread entails a connectivity strength of *g* = 1.5.

**Figure S5.**
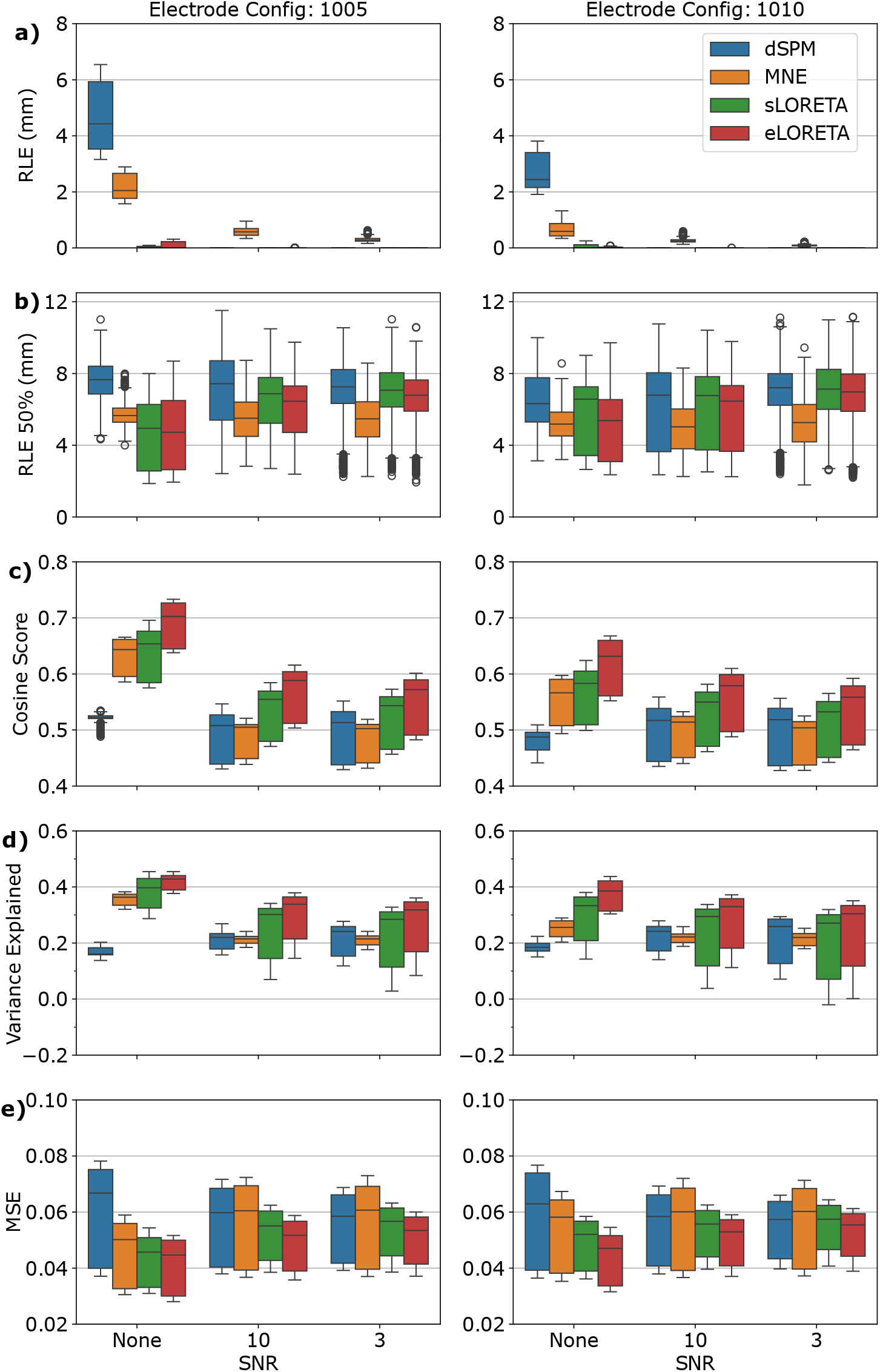
MN-family metrics for loose orientation. Each algorithm from the MN-family is represented by a different colour: dSPM (blue), MNE (orange), sLORETA (green), and eLORETA (red). Rows a–e display the different metrics, different coloured box and whiskers depict the different algorithms, and the x-ticks represent the signal-to-noise ratio in dB. Metrics are computed with a minimally ‘loose’ orientation, meaning that the dipole orientations were allowed to deviate by a factor of up to 5% from the normal of the cortical surface. Even with such a small loose factor, all metrics improved substantially.

**Figure S6.**
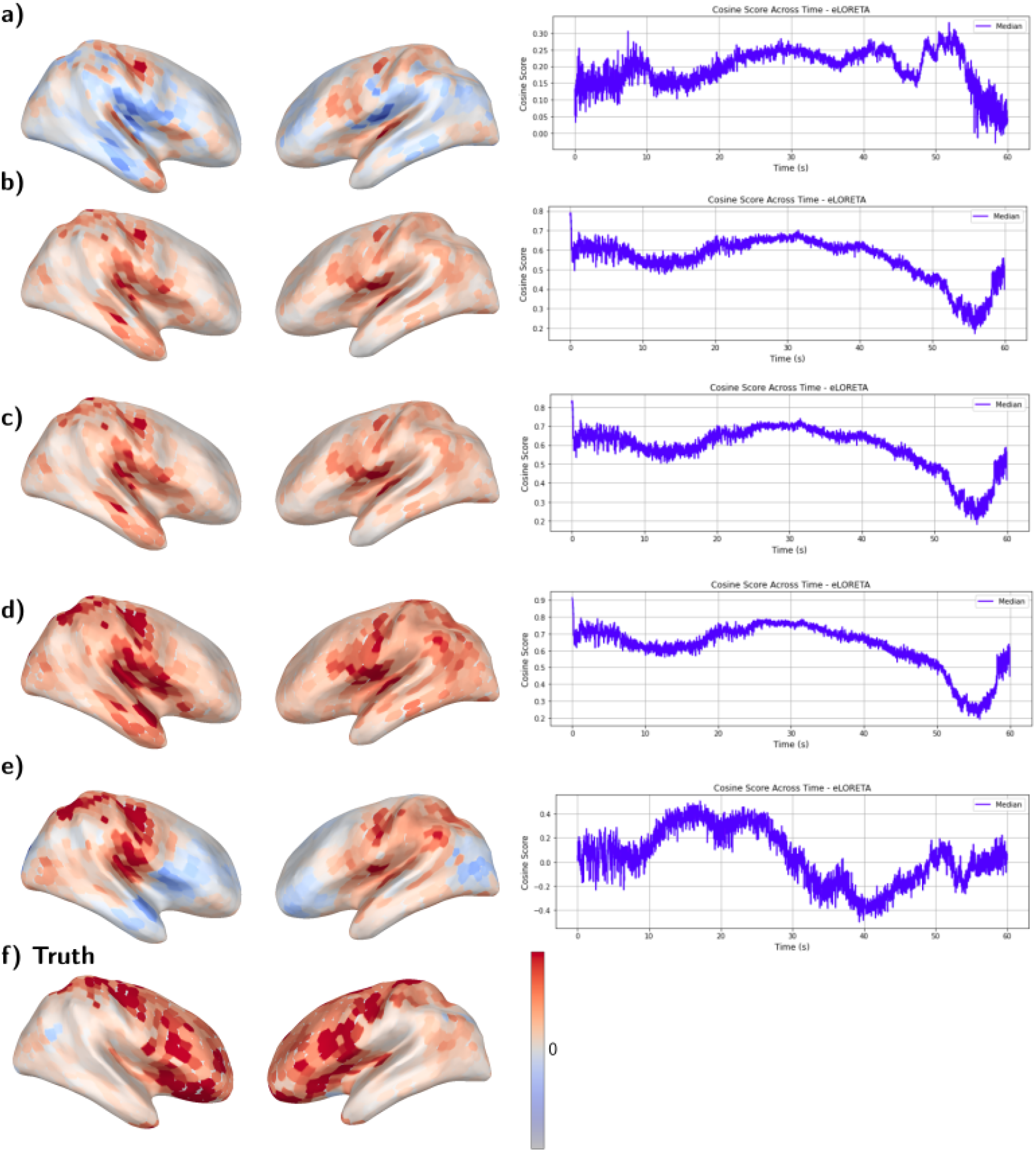
eLORETA method adjusting the loose parameter of a thalamic seizure with SNR 10. The localisation result at *t* = 20 s is visualised on the brain surface. The cosine score across the entire simulation is plotted beside each localisation result. a) Loose = 0 (fixed orientation). b) Loose = 0, with np.abs (estimate). c) Loose = 0.05 (loose orientation). d) Loose = 1 (free orientation). e) Loose = 1 (free orientation), but taking only the normal to the surface component. f) Truth.

**Figure S7.**
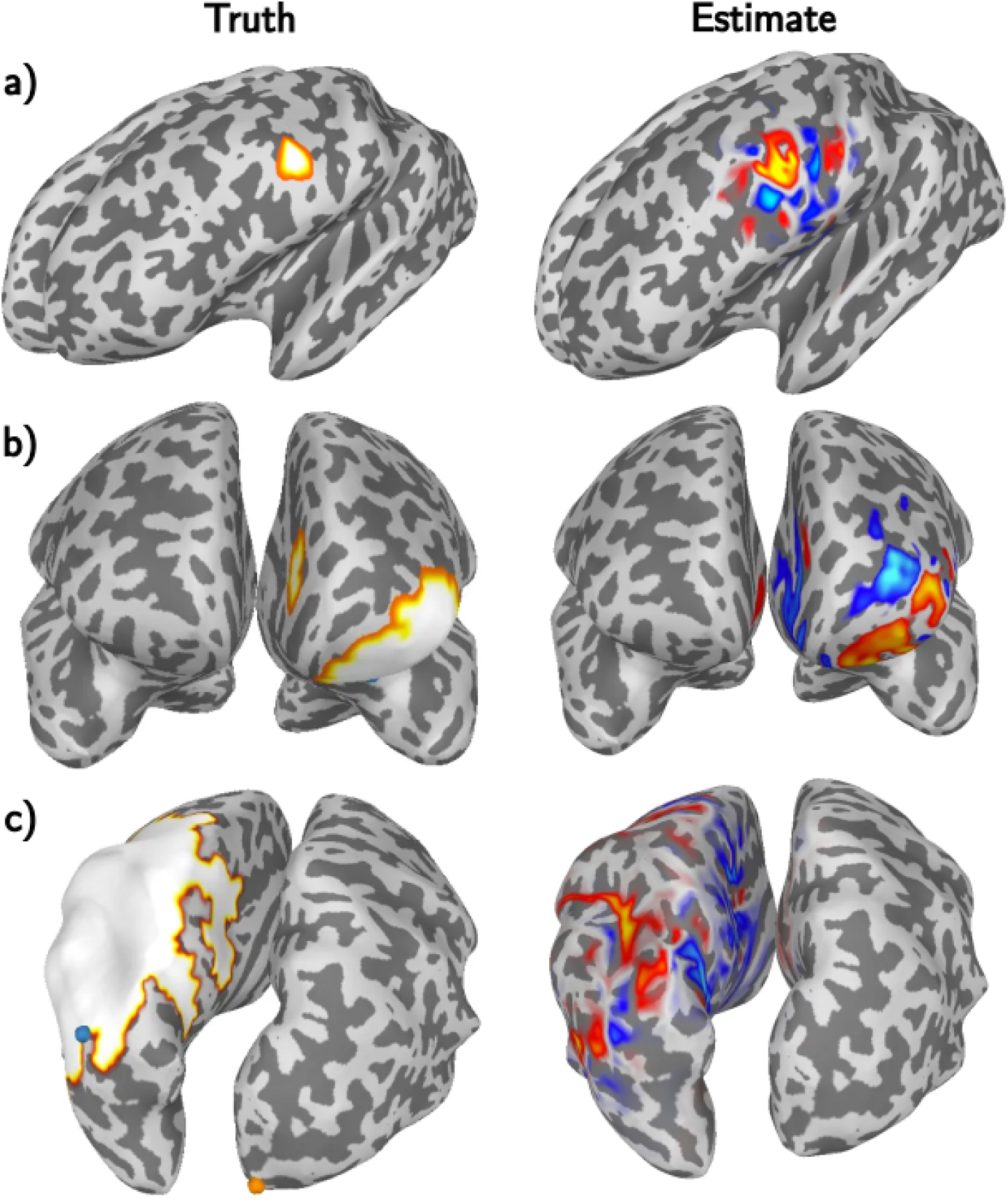
Source estimate against truth for patch sources. The left column is the underlying true source; the right column is the estimate from eLoreta. For the true patch source, all dipoles have the same positive magnitude. For the estimate, the warm colours signify a positive deviation, and the cool colours signify a negative deviation. a) Cortical source of radius 5 mm centred at ‘LH-DorsAttn-PrCv-1’. b) Cortical source of radius 20 mm centred at ‘LH-Cont-PFCv-3’. c) Cortical source of radius 40 mm centred at ‘LH-Cont-IPL-3’.

**Figure S8.**
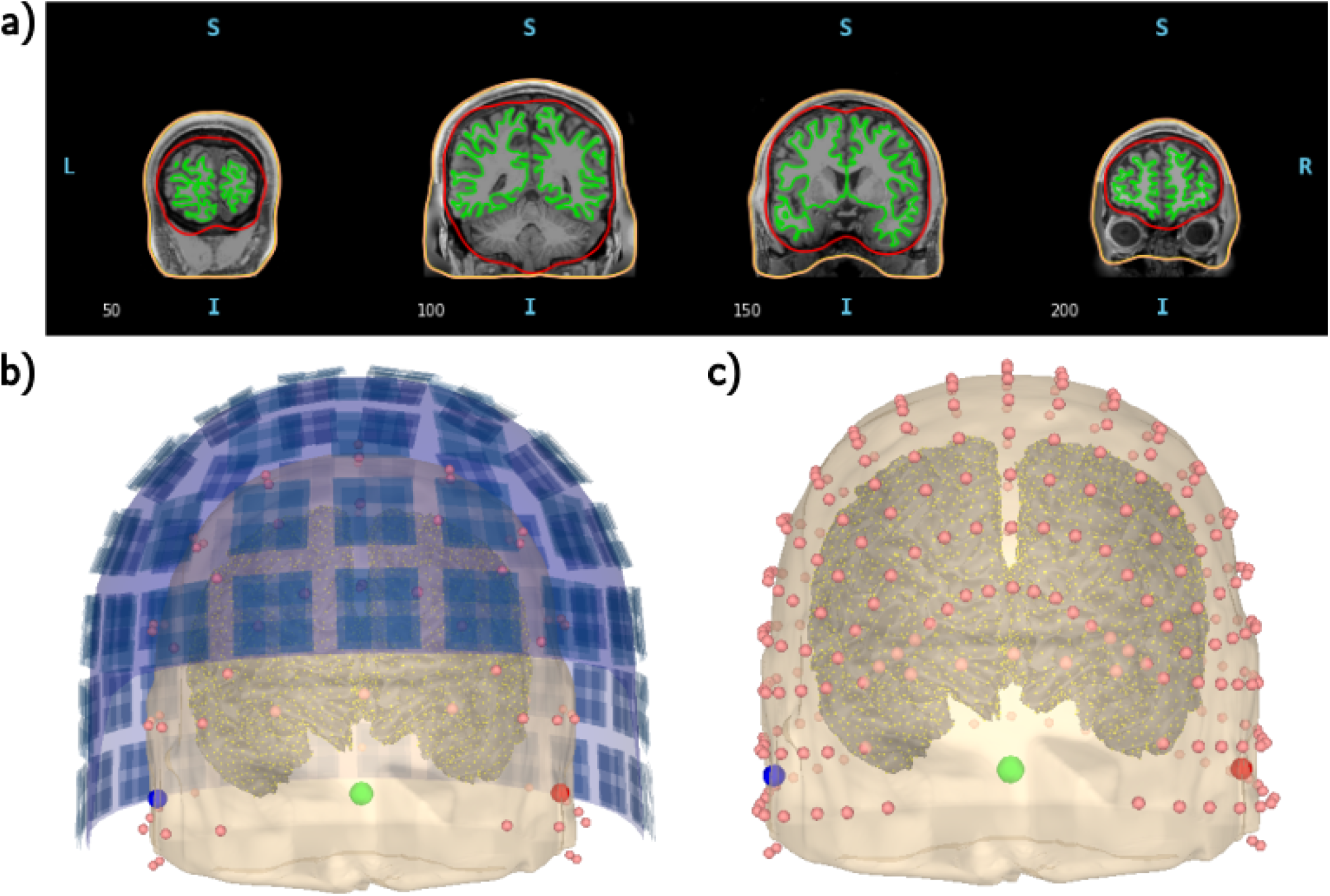
Visualisation of patient MRI surfaces, BEM model, dipole source space, and sensors. (a) The three layers of the BEM model: skin–skull (orange), skull–brain (red), and the white–grey matter boundary (green). (b) MEG sensors are shown as dark blue squares and EEG electrodes (10–10 system) as pink dots. Green, blue and red spheres mark the fiducial landmarks (nasion, right preauricular point, and left preauricular point, respectively). Yellow dots on the cortical surface indicate the dipole locations comprising the source space. (c) EEG electrodes displayed separately using the 10–05 system, without MEG sensors.

