## Supplementary material for "A Neural Mass Modelling Framework for Evaluating EEG Source Localisation of Seizure Activity"

### S1 Supplementary Material

#### S1.1 Epileptor Model Mechanisms

Fig. S1a displays the three subsystems of the Epiletor model that each operate at a different timescale. The fast subsystem ($x_1$, $y_1$ - blue) is based on a modified Hindmarsh–Rose model (Hindmarsh & Rose, 1984) and is responsible for the fast oscillatory activity observed during seizures. This subsystem exhibits bistability between a stable fixed point (representing the interictal, resting state) and a stable limit cycle (oscillatory activity representing the ictal state).

The fast subsystem is coupled to a slow subsystem ($z$ - green), which governs transitions into and out of the ictal state through a hysteresis loop under appropriate parameter regimes (Saggio & Jirsa, 2024). The phase diagram of Fig. S1b displays this hysteresis loop as a clockwise trajectory, forming the core mechanism of the Epileptor model.

Finally, the intermediate-timescale subsystem ($x_2$, $y_2$ - orange) operates near a saddle-node on an invariant circle (SNIC) bifurcation (Houssaini et al., 2020). Each crossing of this bifurcation point by the slow subsystem generates a distinct spike through following an invariant circle trajectory. During the preictal period, these preictal spikes reflect increased excitability of the intermediate subsystem near seizure onset. During the ictal period, the intermediate variables modulate the fast variables such that together they produce the characteristic spike–wave complex ($x_1 - x_2$ in Fig. **??**b).

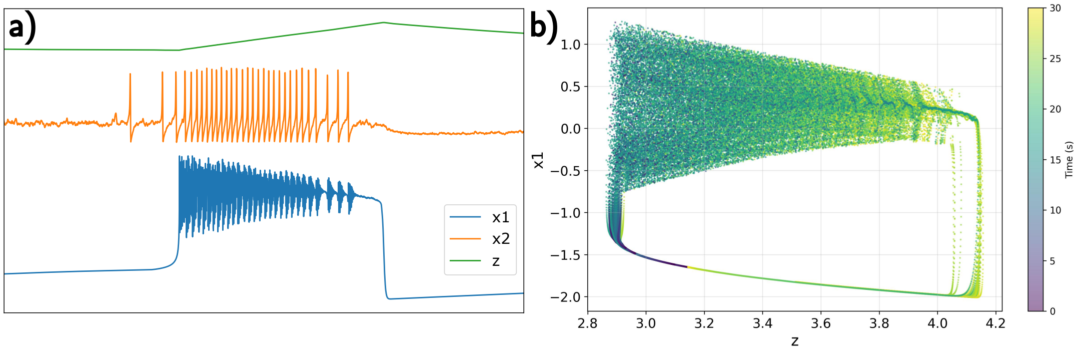


**Figure S1: Mechanisms of the Epileptor model.** a) The Epileptor model can be viewed as the interaction among three subsystems operating at different timescales: $x_1$ (blue) at the fast timescale, $x_2$ (orange) at the intermediate timescale, and $z$ (green) at the slow timescale. b) $x_1 - z$ phase diagram where the colour represents the time of the trajectory. We see that, for a system initialised at $(x_1, z) = (-1.6, 3.1)$, it starts on a fixed point and then moves clockwise in the phase diagram (seen by the colour change from purple to dark green) to reach the unstable limit cycle on the upper portion of the loop. The system then continues along the hysteresis loop until it reaches the fixed point, with the cycle continuing if the simulation was continued.

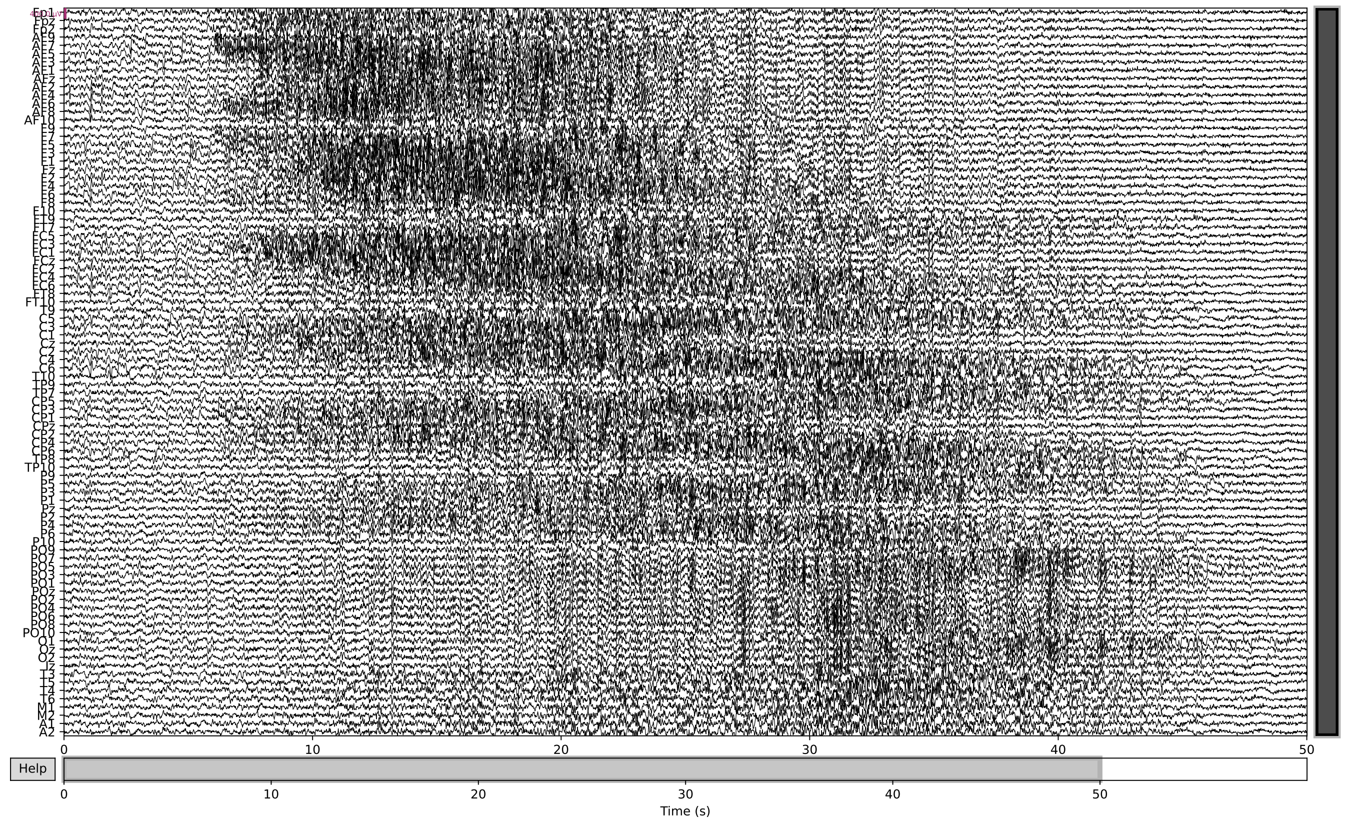


**Figure S2: EEG trace of simulation with SNR of 10 dB.** Each row represents the simulated EEG activity at each labelled electrode. Increase volatility can be seen in areas undergoing epileptic activity. The simulated EEG exhibited clinically consistent features, including focal onset, spatial spread, and temporal evolution with characteristic slowing.

#### S1.2 Analysis of the EEG Activity Generated by the Epileptor Model

As with all simulations, due diligence must be performed to ensure that the simulations meet our requirements. Fig. S2 shows the EEG trace of the sample seizure originating from the thalamus. A neurologist reviewer confirmed that the simulated EEG exhibited clinically consistent features, including focal onset, spatial spread, and temporal evolution with characteristic slowing. Fig. S3 visualises the spectral characteristics of the sample simulation. From Fig. S3a depicts the spectrogram for a single electrode, demonstrating how most power is concentrated at the very low frequencies, although some patches of power are also evident at higher frequencies. In terms of temporal information, it is clear when the seizure is occurring. In Fig. S3b, taking the log of the power reveals two major deviations from the brain's natural $1/f^{\alpha}$ power scaling laws (Donoghue et al., 2020) in the low-frequency and the 15-20 Hz range. These peaks coincide with the 'DC shift' and the 'fast oscillatory activity' of population $x_1$ in the Epileptor model.

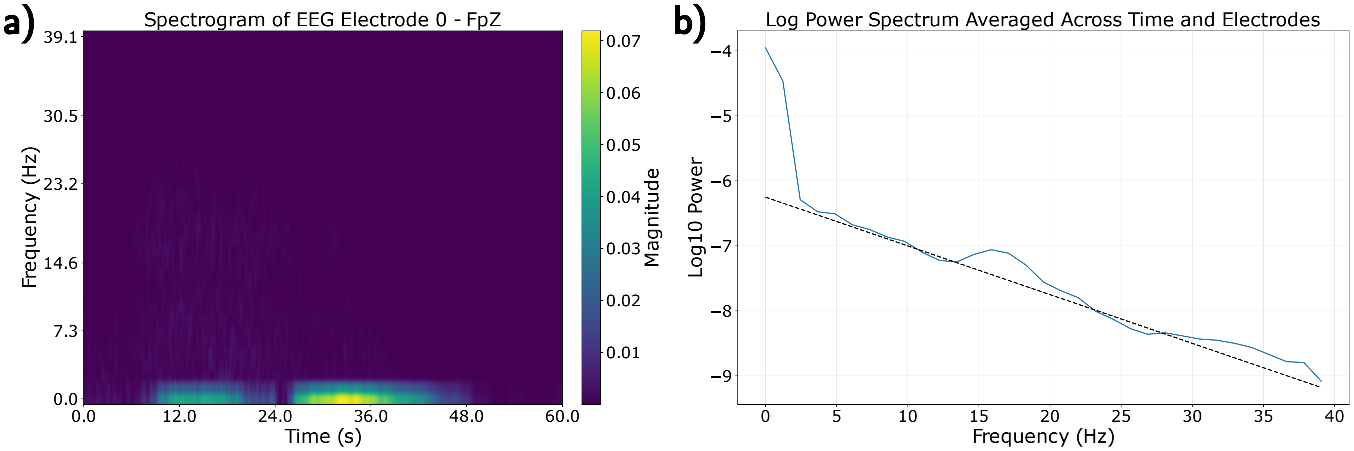


**Figure S3: EEG simulation frequency spectrum.** Spectral properties for the thalamic seizure simulation are graphed. a) Spectrogram for EEG electrode 0 - Fpz across time and frequency. b) Log power spectrum averaged across time and electrodes. The dotted line represents a $1/f^{\alpha}$ power scale ($\alpha = 3/40$ in this instance).

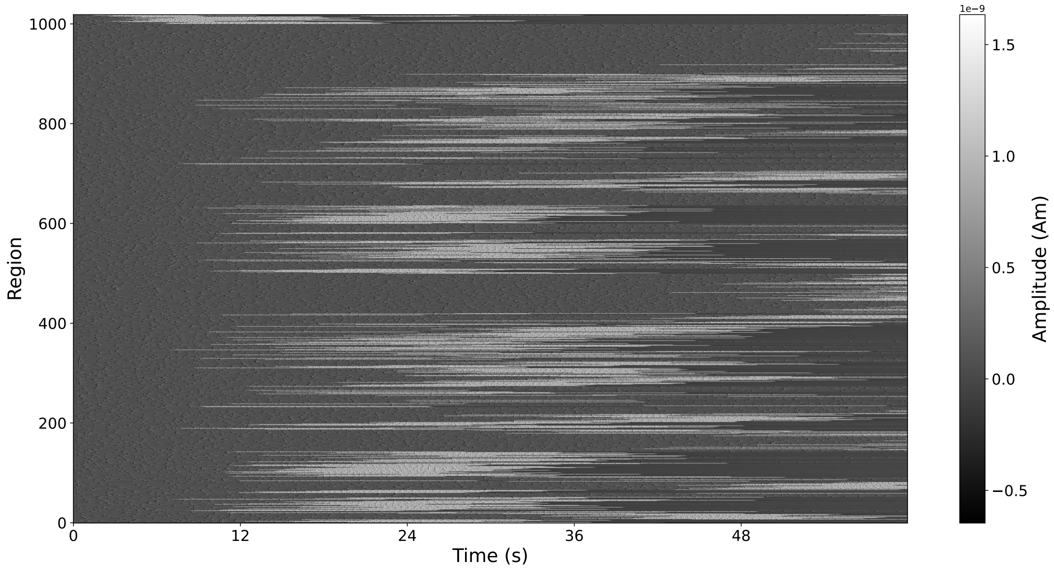


**Figure S4: Carpet plot of seizure originating from thalamus with partial spread.** Partial spread entails a connectivity strength of $g = 1.5$.

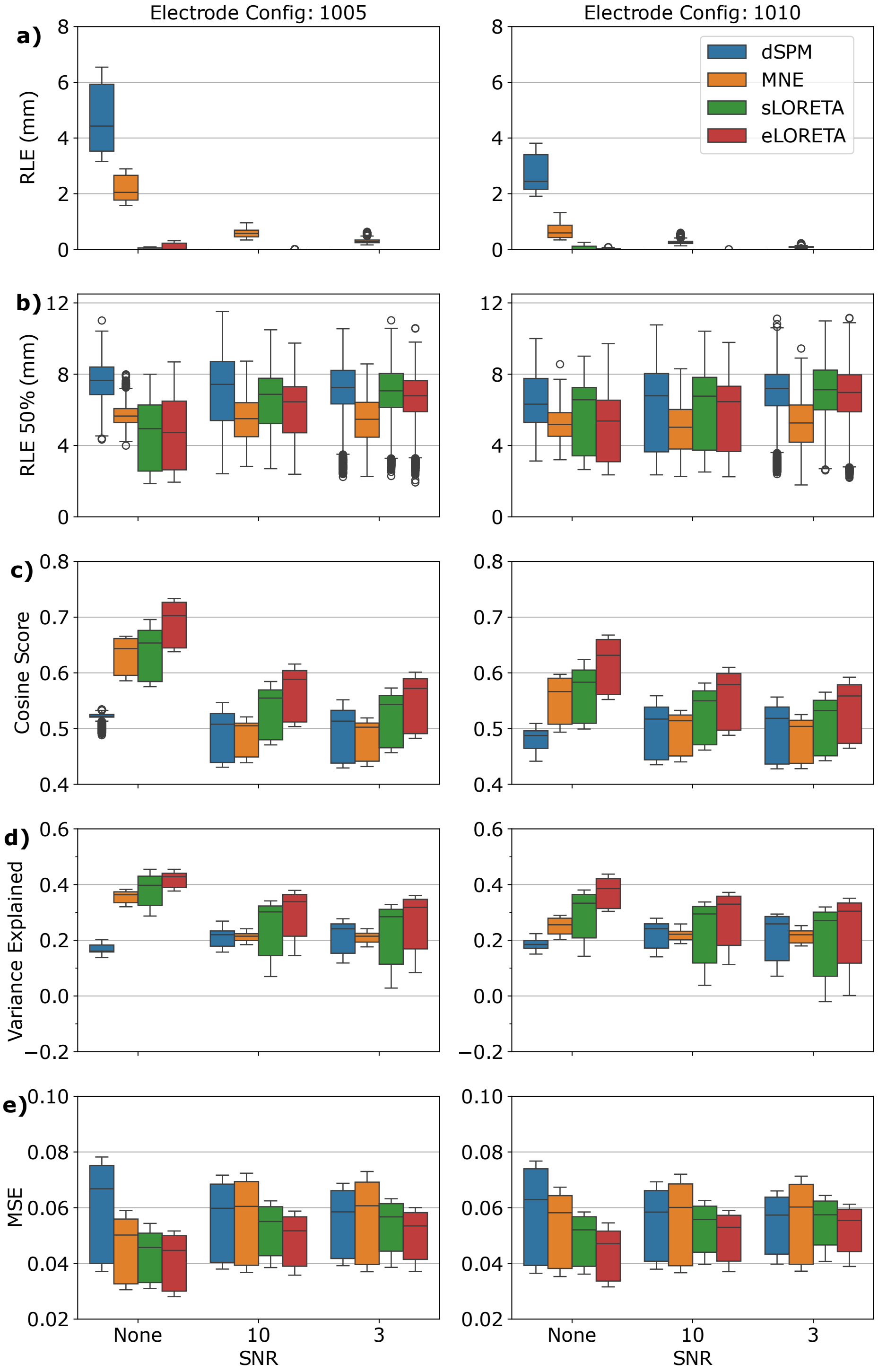


**Figure S5: MN-family metrics for loose orientation.** Each algorithm from the MN-family is represented by a different colour: dSPM (blue), MNE (orange), sLORETA (green), and eLORETA (red). Rows a–e display the different metrics, different coloured box and whiskers depict the different algorithms, and the x-ticks represent the signal-to-noise ratio in dB. Metrics are computed with a minimally 'loose' orientation, meaning that the dipole orientations were allowed to deviate by a factor of up to 5% from the normal of the cortical surface. Even with such a small loose factor, all metrics improved substantially.

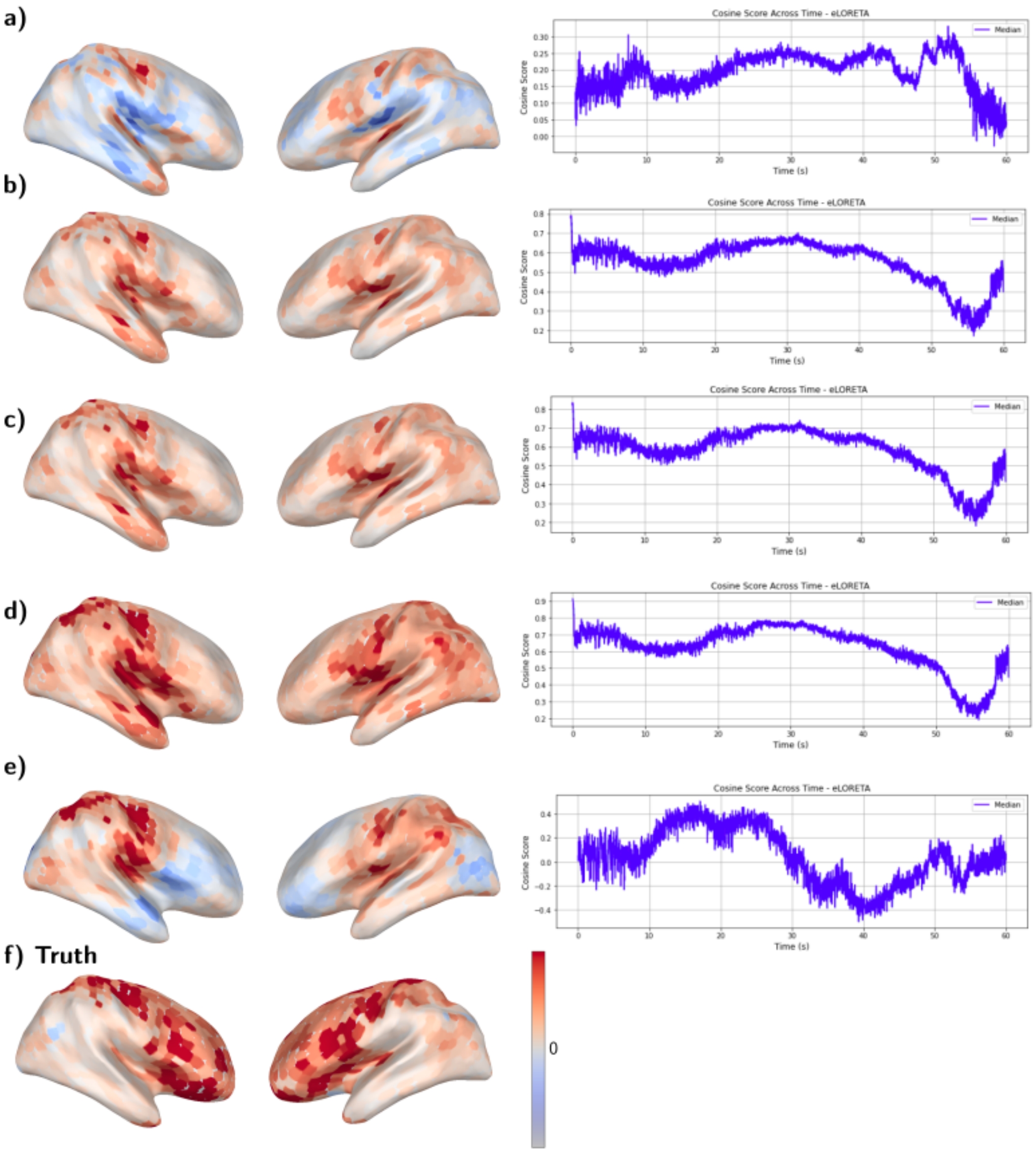


**Figure S6: eLORETA method adjusting the loose parameter of a thalamic seizure with SNR 10.** The localisation result at $t = 20$ s is visualised on the brain surface. The cosine score across the entire simulation is plotted beside each localisation result. a) Loose = 0 (fixed orientation). b) Loose = 0, with np.abs (estimate). c) Loose = 0.05 (loose orientation). d) Loose = 1 (free orientation). e) Loose = 1 (free orientation), but taking only the normal to the surface component. f) Truth.

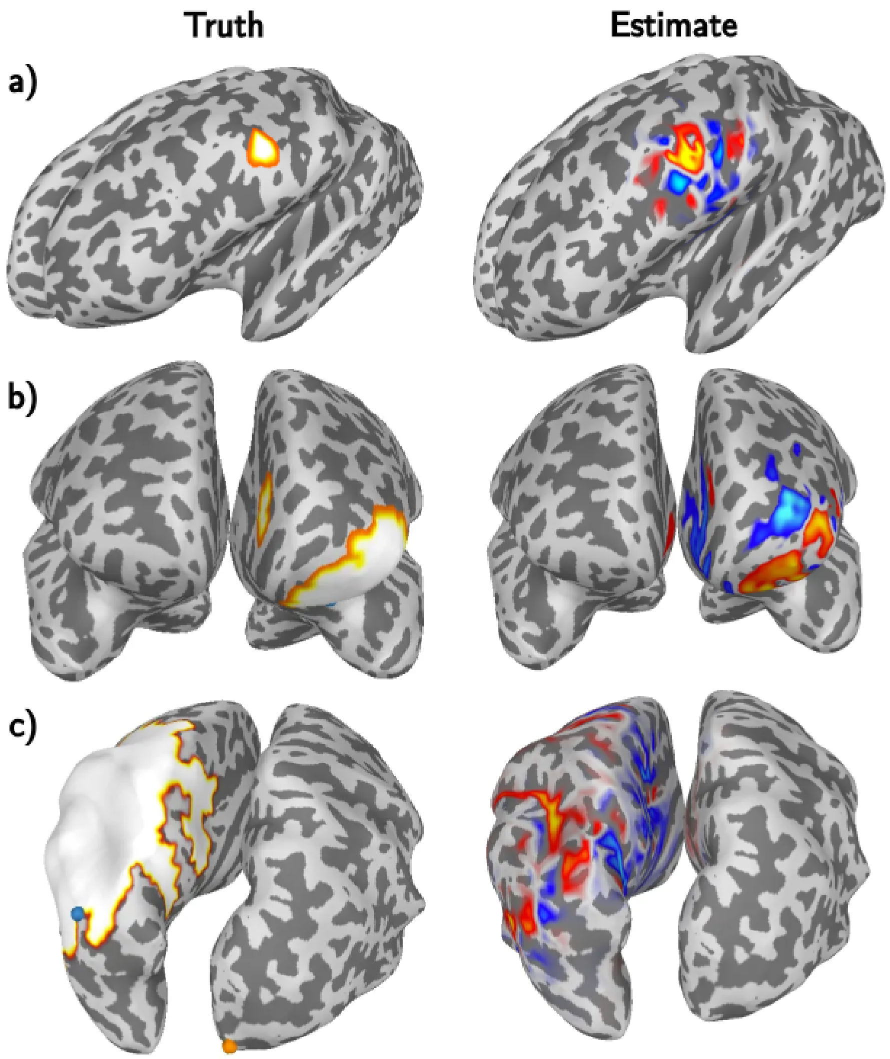


**Figure S7: Source estimate against truth for patch sources.** The left column is the underlying true source; the right column is the estimate from eLoreta. For the true patch source, all dipoles have the same positive magnitude. For the estimate, the warm colours signify a positive deviation, and the cool colours signify a negative deviation. a) Cortical source of radius 5 mm centred at 'LH-DorsAttn-PrCv-1'. b) Cortical source of radius 20 mm centred at 'LH-Cont-PFCv-3'. c) Cortical source of radius 40 mm centred at 'LH-Cont-IPL-3'.

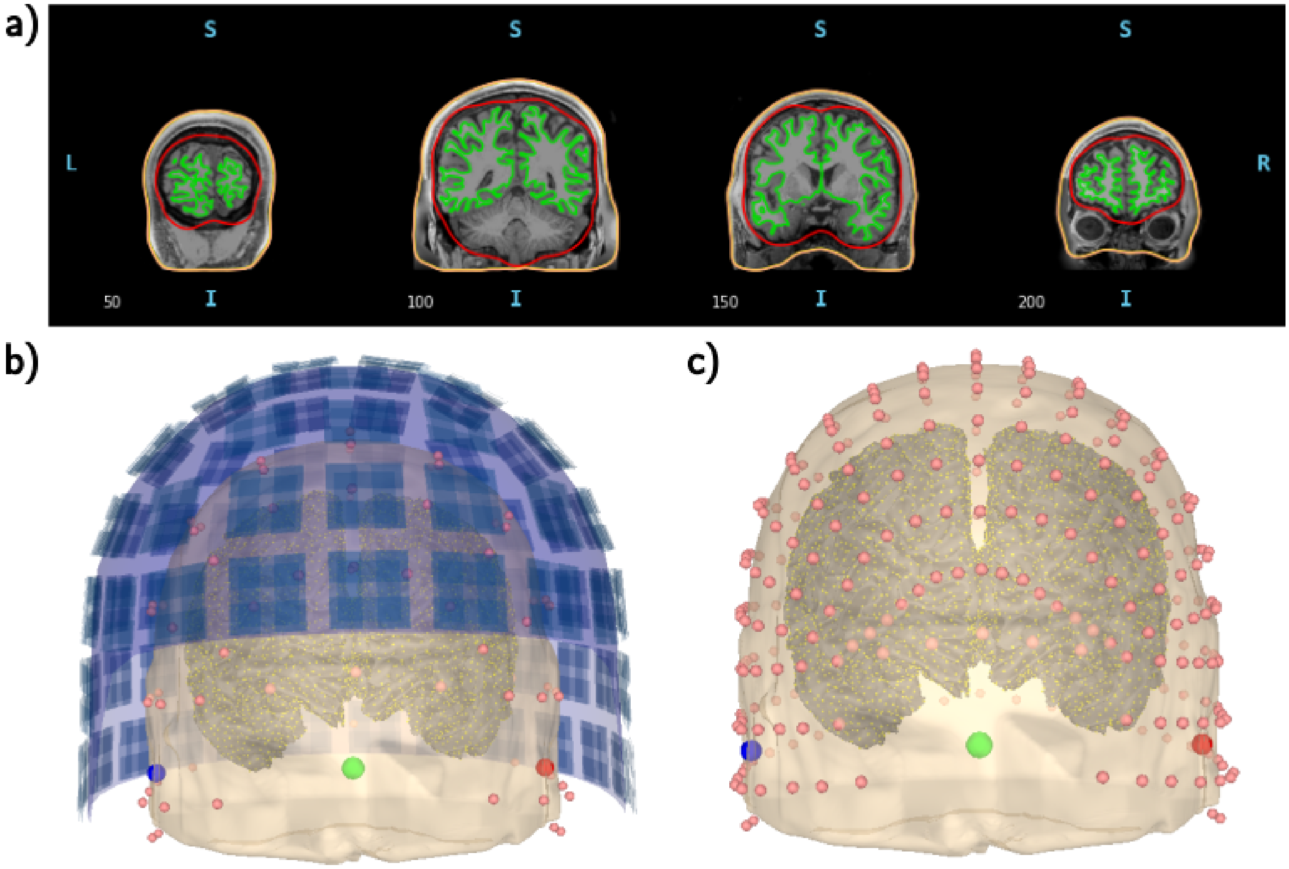


**Figure S8: Visualisation of patient MRI surfaces, BEM model, dipole source space, and sensors.** (a) The three layers of the BEM model: skin–skull (orange), skull–brain (red), and the white–grey matter boundary (green). (b) MEG sensors are shown as dark blue squares and EEG electrodes (10–10 system) as pink dots. Green, blue and red spheres mark the fiducial landmarks (nasion, right preauricular point, and left preauricular point, respectively). Yellow dots on the cortical surface indicate the dipole locations comprising the source space. (c) EEG electrodes displayed separately using the 10–05 system, without MEG sensors.
